# Synthetic lethality of p97 inhibition and PSMC2-overexpression overcomes bortezomib resistance in multiple myeloma

**DOI:** 10.64898/2026.11.02.761087

**Authors:** James J. Ignatz-Hoover, E. Ricky Chan, Ehsan Malek, John D’Cruz, Nida Mubin, Wei Huang, Derek J. Taylor, Mehmet K. Samur, Nikhil C. Munshi, Kenneth C. Anderson, James J. Driscoll

**Affiliations:** Division of Hematology & Oncology, Case Western Reserve University, Cleveland, OH USA; Adult Hematologic Malignancies & Stem Cell Transplant Section, Seidman Cancer Center, University Hospitals Cleveland Medical Center, Cleveland, OH USA; Case Comprehensive Cancer Center, Case Western Reserve University, Cleveland, OH USA; Cleveland Institute for Computational Biology, Case Western Reserve University, Cleveland, OH USA; Roswell Park Comprehensive Cancer Center, Buffalo, NY USA; University School, Hunting Valley, OH USA; Case Western Reserve University Scientific Enrichment and Opportunity/Youth Engaged in Science Program, Cleveland, OH USA; Departments of Biochemistry and Pharmacology, Case Western Reserve University, Cleveland, OH USA; Department of Biostatistics and Computational Biology, Dana-Farber Cancer Institute and Harvard School of Public Health, Boston, MA, USA; Jerome Lipper Multiple Myeloma Center, LeBow Institute for Myeloma Therapeutics, Dana-Farber Cancer Institute, Harvard Medical School, Boston, MA, USA

**Keywords:** Multiple myeloma, proteasome, PSMC2 AAA-ATPase, p97, synthetic lethality, chemoresistance

## Abstract

The aim of the present study was to analyze genomic profiles obtained from multiple myeloma (MM) patient tumors to identify prognostic classifiers of clinical response and survival following bortezomib monotherapy and then target key actionable classifiers that underly bortezomib resistance. We discovered higher levels of proteasome *PSMC2* messenger RNA in myeloma cells from patients that did not respond to bortezomib and that higher *PSMC2* expression correlated with inferior progression-free and overall survival. Pharmacologic screening revealed that CB-5339, an ATP-competitive inhibitor of the molecular chaperone p97/VCP that shuttles ubiquitin-conjugated substrates to proteasomes, was synthetic lethal in MM cells engineered to overexpress *PSMC2*. p97 co-immunoprecipitated with 26S proteasomes affinity-purified from *PSMC2-*overexpressing cells which exhibited greater activity and reduced bortezomib sensitivity relative to *PSMC2-KO* cells. Bortezomib treatment followed by *in-situ* cross-linking and western blotting revealed higher levels of K48- and K63-linkage-specific ubiquitin chains covalently bound to 26S proteasomes isolated from *PSMC2*-*KO* cells compared to proteasomes affinity-purified from *PSMC2*-overexpressing cells. Western blotting demonstrated higher levels of 19S RPs, 26S proteasomes, and a PSMC2-PSMC1-PSMD2 subcomplex in *PSMC2*-overexpressing cells relative to *PSMC2*-*KO* cells. CB-5339 and bortezomib co-treatment synergistically targeted tumor cells obtained from MM patients previously exposed to bortezomib therapy and reduced the *in vivo* growth of *PSMC2*-overexpressing myeloma xenografts in mice. Taken together, we have discovered a genetic vulnerability that provokes proteasome-intrinsic bortezomib resistance and validated the potential of synthetic lethality to guide therapeutic interventions. Our study reaffirms that transcriptomic heterogeneity in MM is a primary driver of prognostic variability.

**Key points:** *PSMC2* overexpression in MM cells promotes association of p97 with proteasomes that demonstrate higher activity and bortezomib resistance.

*PSMC2* overexpression is synthetic lethal with pharmacologic inhibition of p97 ATPase activity and overcomes bortezomib resistance *in vivo*.

**Novelty.:** Higher expression of the proteasome ATPase *PSMC2* in tumor cells from MM patients correlated with reduced clinical response to bortezomib and inferior PFS and OS. *PSMC2* overexpression is synthetic lethal combined with pharmacologic inhibition of p97 ATPase activity and overcomes bortezomib resistance. We discovered a genetic event that provokes proteasome-intrinsic adaptations but also represents an actionable vulnerability that can be exploited using synthetic lethal precision medicines.

**Statement of Significance.:** Synthetic lethality is a transformative strategy in precision medicine to exploit otherwise undruggable targets and eliminate disease-causing cells while sparing healthy tissue. CB-5339 is an ATP-competitive p97 inhibitor that triggered synthetic lethality in *PSMC2*-overexpressing cells. PSMC2 promotes physical association of the molecular chaperone p97 with proteasomes and confers a previously unrecognized mechanism of proteasome-intrinsic bortezomib-resistance. Pharmacologic inhibition of p97 ATPase activity is synthetic lethal in *PSMC2*-addicted cells *in vitro* and *in vivo*. We have identified an actionable, cancer-specific vulnerability with translational avenues to tailor proteasome and p97-directed strategies that mitigate relapse risk and improve patient outcomes.

## Introduction

Multiple myeloma (MM) treatment presents challenges due to genetic heterogeneity, complex pathophysiology and the lack of unifying pathognomonic events^1–5^. Proteasome inhibitors (PIs) have extended patient survival, however, those with certain cytogenetic and clinical features are at risk of poor outcomes^6–8^. Management of relapsed and/or refractory (RR)MM is a vital aspect of patient care and deciphering the conundrum of treatment resistance remains unmet^9–11^. Since MM cells are antibody-producing factories, one strategy is to exploit their reliance on proteostasis^12–14^.

Proteasomes are structurally and functionally heterogeneous macromolecules that regulate the proteome through selective and efficient protein degradation^15,16^. Bortezomib (Btz) reversibly inhibits the chymotrypsin-like (ChT-like) activity at the PSMB5-subunit located in the proteasome 20S core particle (CP). 20S CPs are capped by 19S regulatory particles (RPs) which consist of heterohexameric AAA-ATPases encoded by *PSMC1-6* and lids encoded by *PSMD* genes^17,18^. PSMC2 is a AAA-ATPase responsible for unfolding and translocating substrates through 19S RPs that are threaded into 20S CPs^19–23^. Prior studies reported non-equivalent functions for individual *PSMC* family members^21–26^. Since many patients do not respond to therapy, we sought to identify proteasome-intrinsic vulnerabilities through an integrated analysis of gene expression profiles and clinical datasets. Using this pipeline, we discovered that greater *PSMC2* expression correlated with reduced clinical response to Btz and inferior survival.

## Methods

### Cell lines

MMCLs (ATCC) were cultured in RPMI-1640, and HEK293T (ATCC) was cultured in DMEM. Media was supplemented with 10% Hyclone FBS and penicillin/streptomycin.

### Western blots

Cells were washed, resuspended in RIPA or Cel-Lytic M buffer, centrifuged at 12,000 x g for 20 min and protein quantified. Samples were electrophoresed on 4-12% gels and probed with antibodies to PSMC1, PSMC2, PSMC6, K27-, K48- and K63-linkage-specific Ub (Abcam), PSMB5, p97, PSMD14, and GAPDH (Cell Signaling Technologies), FLAG-M2 (Sigma), ECM29, ZFAND1, K11- and K29-linkage-specific Ub antibodies (Thermo-Fisher), PSMB8 (Santa Cruz Biotechnology), K6- and K33-linkage-specific Ub (Antibodies.com).

### Vectors

*Myc-DDK-PSMC2/6* and VCP-GFP vectors (Origene), *pLKO* plasmid (Sigma-Aldrich), lenti-Luc-Neo and PSMD14-TEV-glutathione S-transferase (GST) (Addgene) and Cherry two-plasmid inducible p97-KD system (VectorBuilder). The *VB240429-1448qhf* mCherry element expressed a constitutively active *TET*-inducer. mCherry positive cells were sorted with a FACS-ARIA-SORP (BD Biosciences). BSD-TRE3G expresses GFP and 3 shRNAs that target *p97* under TRE3G control (5’-GAATAGAGTTGTTCGGAATAA-3’, 5’-ACCGTCCCAAT CGGTTAATTG-3’; and 5’-ATATAGAGAGTAGGTTGATTT-3’).

### Chemicals

Reagent grade Btz, carfilzomib (Cfz) (Selleck Chemicals), CB-5339 (MedChemExpress), blasticidin, neomycin, and puromycin (Thermo-Fisher) were used.

### *CRISPR/CAS9*-mediated gene knockout

Cells were transfected using Neon electroporation *per* manufacturer’s protocol **using g**uide RNAs, CAS9 enzyme, and scrambled controls (5’-UGAUCGGCACCGAGGUAAUC-3’, 5’-CAUGUCAACUGACCUCGGAU-3’, 5’-AGGAGCUGGAUUCCCGCUCC-3’) (Synthego).

### Lentiviral transduction and selection

Vectors were co-transfected with helper plasmids pRSV-REV, pMDLg/pRRE, and pCMV-VSV-g in HEK293T cells with Turbofectin and cultured in OPTiMEM. After transfection, supernatant was filtered, added to target cells with polybrene (Sigma Aldrich), spun at 800 x g, cultured for 18 h, and polybrene removed. Cells were selected in blasticidin (10 μg/mL), puromycin (2 μg/mL), and neomycin (250 µg/ml). **Immunoprecipitation.** Lysates were prepared using Cel-Lytic M supplemented with 1 mM ATP, 10% glycerol, 1 mM DTT, and 1 mM MgCl_2_. Protein was quantified using Pierce Protein-Plus. Proteins were immunoprecipitated using GST, FLAG, or Magstrep beads according to manufacturer’s instructions. Glutathione and Twin-Strep constructs were digested with TEV protease for 2 h at RT. FLAG constructs were eluted using 3X-FLAG peptide (Sigma).

### Crosslinking of Ub-chain-specific conjugates to affinity-purified proteasomes

Cells were crosslinked using dithiobis(succinimidyl propionate, DSP) and resuspended in anhydrous DMSO. Cells were pretreated with Btz (10 nM) for 18 h, and washed with cold PBS. DSP was diluted into cold PBS to a final concentration of 1 mM and applied to cells for 60 min at 4 °C with gentle rocking. Crosslinking was quenched by addition of 25 mM Tris-HCl (pH 7.5) for 10 min at RT, followed by a PBS wash. Cells were lysed on ice in 1% Triton X-100, HEPES-NaCl pH 7.4 supplemented with protease inhibitors and 10 mM N-ethylmaleimide, 1 mM ATP, 1 mM DTT, 1 mM MgCl_2_ and 10% glycerol, and clarified by centrifugation. Lysate was immunoprecipitated using glutathione-sepharose beads, washed, protein eluted, and probed by western blotting.

### Proteasome activity

Proteasomes were plated in 20 mM Tris-HCl, pH 7.6, 1 mM ATP, 1 mM DTT, 1 mM MgCl_2,_ and Suc-LLVY-AMC (100 uM) and fluorescence recorded.

### Viability assays

XTT and Alamar Blue dye were used to measure cell viability.

### Animal studies

Animal studies were approved by Institutional Animal Care and Use Committees. Sixty-four female, NOD-SCID-γ mice (Jackson Lab) were injected sc with 1.5x10 ARH-77-*pLKO-LUC* or ARH-77-*PSMC2-FLAG-LUC* cells in the right flank. After 7 days, tumors were assessed by BLI (IVIS Spectrum).

### Biostatistics

Data was analyzed using Graphpad. Co-occurrence analysis was performed using Fisher’s exact test to generate an Odds ratio and *p*-value, n given in each case. Odds ratio was calculated for treatment response using Epitools package (Comprehensive R Archive network, CRAN) using 95% CIs and Fisher’s exact test. OS including Kaplan-Meier (K-M) curves and Cox proportional hazard regression for expression tertile comparisons was performed using the Survival package (CRAN) and significant differences were identified using a Chi-square test. Bioassays represent three biological replicates unless otherwise stated. Error bars represent +/-1 SD, except in the mouse data where they represent +/-1 SE of the mean. Dose curves were analyzed using nonlinear regression analysis. R^2^ values and resultant LD_50s_ are reported. The difference between two means was compared using two-sided, unpaired Student’s t-test. Sidak’s multi-comparison testing compared differences between multiple measurements of two groups and

Dunnett’s multi-comparison testing compared differences between multiple samples in ≥ 3 groups.

### Bliss independence synergy analysis

Drug interactions were evaluated using the Bliss independence model^27^. The fraction affected (Fa) for each condition was calculated as Fa = 1 − (viability − blank)/(vehicle − blank), averaged across technical replicates (n = 3). For each combination well, the Bliss-expected Fa was calculated from the single-agent Fa values measured on the same plate as: Fa_expected = Fa_BTZ + Fa_CB − (Fa_BTZ × Fa_CB). The Bliss excess (synergy score) for each combination was then defined as the difference between the observed and expected Fa: Bliss excess = Fa_observed − Fa_expected. Positive Bliss excess values indicate synergy (greater-than-additive effect). Calculations were performed in Python (NumPy).

## Results

### *PSMC2*-overexpression correlates with reduced patient response to Btz and inferior survival

We hypothesized that expression of individual proteasome genes could inform clinical responses to Btz-based therapies in MM patients. The pivotal phase III APEX trial (NCT00073280) randomized MM patients to either Btz or high-dose dexamethasone (Dex) monotherapy. Gene expression profiling data from these patients, generated on Affymetrix microarrays (GSE9782), were used to identify predictive biomarkers, molecular signatures of disease progression, and map patient heterogeneity (Fig. S1). We compared the expression of individual proteasome genes in myeloma cells isolated from patients who were either Btz responders (Btz-Rs) or non-responders (Btz-NRs). We identified twelve genes that were significantly overexpressed in Btz-NRs, but nine of these genes (*PSMA3, PSMB4, PSMB5, PSMC6, PSMD1, PSMD4, PSMD7, PSMD11,* and *PSMD12*) also correlated with clinical response to Dex (Table S1). Higher expression of the proteasome 19S RP AAA-ATPase subunits *PSMC2* and *PSMC6* also correlated with inferior PFS following treatment with Btz, but not Dex (Fig. 1a; Fig. S2, S3). K-M analysis of patients in the highest tertile of *PSMC2* expression correlated with inferior PFS relative to patients in the low/medium tertiles following treatment with Btz, but not Dex (Fig. 1b). Patients in the highest tertile of *PSMC2* expression also correlated with inferior OS relative to patients in the low/medium tertiles (Fig. 1c, d). The effect of *PSMC2* expression on OS was specific to Btz treatment (Fig. 1c; Fig. S4, S5). *PSMC2* was the only proteasome gene in which higher expression correlated with increased risk of inferior PFS and OS following Btz, but not Dex, treatment. *PSMC2* messenger RNA (mRNA) expression was greater in tumor cells from Btz-NRs relative to Btz-Rs, but expression was not statistically different between Dex-Rs and Dex-NRs (Fig. 1e). To corroborate findings from the APEX dataset, we analyzed the Relating Clinical Outcomes in MM to Personal Assessment of Genetic Profile study (CoMMpass) in which patients were treated with standard-of-care combinations. Patients with tumors that expressed *PSMC2* in the highest tertile exhibited inferior PFS and OS compared to those with *PSMC2* expressed in the middle or lowest tertile (Fig. 1f, g, Fig. S6-S8). To further interrogate the prognostic significance of *PSMC2*, we performed Cox multivariate regression analyses of the APEX dataset by analyzing gender, number of treatment lines, age, CRP, β-2 microglobulin, albumin, and genetic risk factors; and analyzed CoMMpass using age, SCT status, ECOG performance, gender and R-ISS stage. Multivariate analyses indicated a significant effect of high *PSMC2* expression on OS for patients on either study (Fig. 1h). High *PSMC2* expression also trended towards significance effect on PFS patients on CoMMpass and Btz-treated on APEX (Fig. S8). The results strongly suggested that *PSMC2* is a significant adverse prognostic marker after adjustment for clinical co-variates, and is among the most prognostically informative proteasome subunits, especially for OS.

**Fig. 1.**
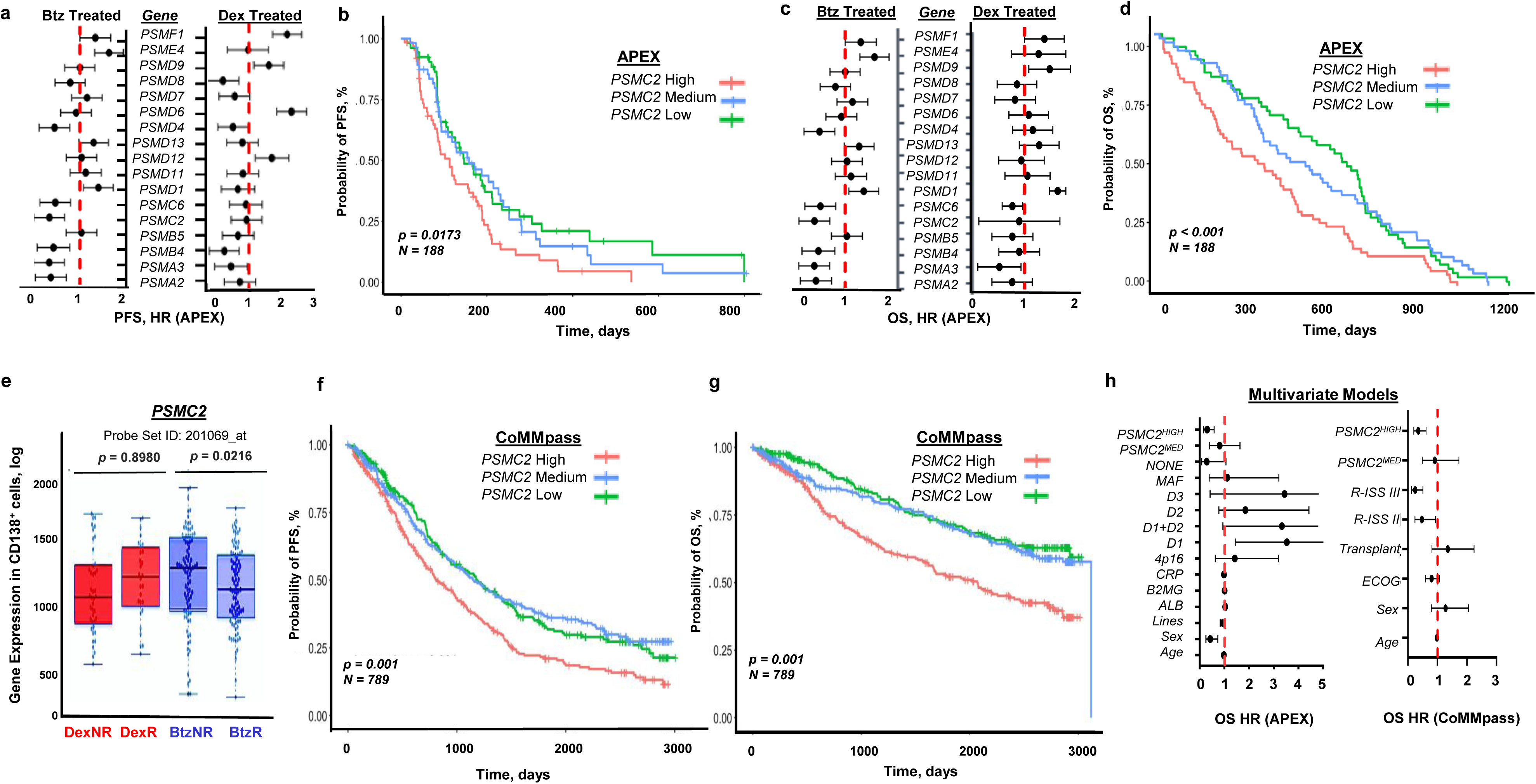
Proteasome gene expression correlates with MM patient response to Btz, OS and PFS. **a**. The APEX trial was a randomized, prospective, institutional review board (IRB)-approved phase III APEX study was conducted at 95 centers worldwide that consented and enrolled 669 patients based upon relevant inclusion and exclusion criteria with RRMM treated with 1-3 lines of prior therapy in accord with the Declaration of Helsinki^62, 63^. Univariate Cox proportional hazard analysis was performed using the APEX data to correlate proteasome gene expression with disease progression for patients treated with either Btz or Dex. Shown are hazard ratios (HRs) for PFS +/-2 standard errors (SE) for low and medium expression tertiles of each gene, relative to the highest expression tertile (reference). Values < 1 therefore indicate that higher expression of that gene is associated with increased risk of disease progression or death. **b**. K-M analysis to compare PFS for patients in the upper tertile of *PSMC2* expression (red) to middle (blue) and lower tertile of expression (green) in patients treated with Btz (n = 188 events). Patients with higher *PSMC2* expression had a PFS of 105 days vs. 155 days for the other two tertiles. **c**. Univariate Cox proportional hazard analysis was performed using the APEX dataset to correlate OS outcomes and gene expression data in patients treated with Btz or Dex. Shown are HRs for OS +/-2 SE for the low and medium expression tertiles of each gene, relative to the highest expression tertile (reference). Values < 1 therefore indicate that higher expression of that gene is associated with increased risk of death. **d**. K-M analysis to compare OS of patients within the high, medium, or low tertile of *PSMC2* expression in patients treated with Btz (n = 188 events). High expression tertiles exhibited statistically significant differences in OS by log-rank test (p = 6x10^-5^; median OS 300 days for the highest tertile vs. 750 days for the lowest). **e**. Box and whisker plot for *PSMC2* RNA expression levels stratified by treatment with Btz or Dex and response status (NR = non-responder, R = responder). Btz R (n = 55) on average exhibited lower levels of *PSMC2* compared to Btz NR (n= 61). Data was analyzed utilizing a two-tailed Student’s t-test; p-values are given within the figure. **f**. The CoMMpass dataset is an international multicenter observational study encompassing >1,100 patients that follows longitudinal molecular and clinical outcomes in MM patients treated with modern induction regimens (Dataset IA22 last accessed June 2026). K-M analysis of the CoMMpass dataset revealed reduced PFS in patients in the high tertile of *PSMC2* expression than those in the low tertile (p-value = 0.001) respectively by the log-rank test, n=263/tertile). **g**. K-M analysis of the CoMMpass dataset revealed worse OS in patients in the high tertile of *PSMC2* expression than those in the low tertile. p-value = 0.001 by the log-rank test, n=263/tertile. **h**. Cox multivariate hazard analysis was performed on patients from the APEX and CoMMpass datasets. The APEX model included biological sex, number of treatment lines, age, CRP, β-2 microglobulin, albumin, genetic risk factors, and PSMC2 expression. The CoMMpass model included age, SCT status, ECOG performance, biological sex, R-ISS stage, and PSMC2 expression. PSMC2 expression is modelled with the lowest tertile as reference, so HR > 1 implies the given variable increases risk of death. Resulting HRs are given +/-SEs.

Project Achilles is a systematic effort to identify genetic vulnerabilities across a panel of genomically characterized cancer cell lines^28,29^. Compared to all cancer cell lines, DNA copy number gain for *PSMC2* was the highest among *PSMC* family members (Fig. S9a). Large-scale functional screening in Project Achilles provided a complementary approach to determine the individual gene effect on cell viability. After *CRISPR/CAS9* knockout (*KO*) or genome-wide shRNA-mediated silencing (RNAi score), the effect on viability among the *PSMC* family was greatest for *PSMC2* (Fig. S9b). The deleterious effect of *PSMC2*-*KO* on cell survival across the Project Achilles panel support the broad and profound impact of this gene.

### *PSMC2* is a unique, independent prognostic factor of Btz resistance

*PSMC2* RNAseq data obtained from MM cells of patients enrolled on the CoMMpass trial revealed a right-sided skew (Fig. S10). The human *PSMC2* gene is located on chromosome **7** at band q22.1-q22.3, and gain of chromosome 7 is a recognized copy number variation in MM. Gain of 7q is associated with hyperdiploid disease which typically involves trisomies of odd-numbered chromosomes and confers a relatively standard-risk prognosis.^30^ We tested if *PSMC2* mRNA levels correlated with 7q gain. Chromosome 7q status was derived from fluorescence *in situ* hybridization (FISH) cytogenetic data and encoded as a binary variable indicating presence or absence of gain. The relationship between *PSMC2* expression and 7q gain was assessed by comparing expression distributions between groups. The results did not reveal a statistical difference in the level of *PSMC2* expression between patients with or without gain 7q (Fig. 2a). We also compared OS outcomes by K-M analysis for patients within each tertile of *PSMC2* expression and 7q status. The results indicated that patients with high *PSMC2* and no 7q gain exhibited worse OS relative to other cohorts (highest vs. low/medium *PSMC2*, +7q gain vs. no 7q gain) (Fig. 2b). The data suggested that *PSMC2*-high expression was not confined to patients with 7q gain, and that *PSMC2*-high expression is not simply a binary surrogate for 7q gain.

**Fig. 2.**
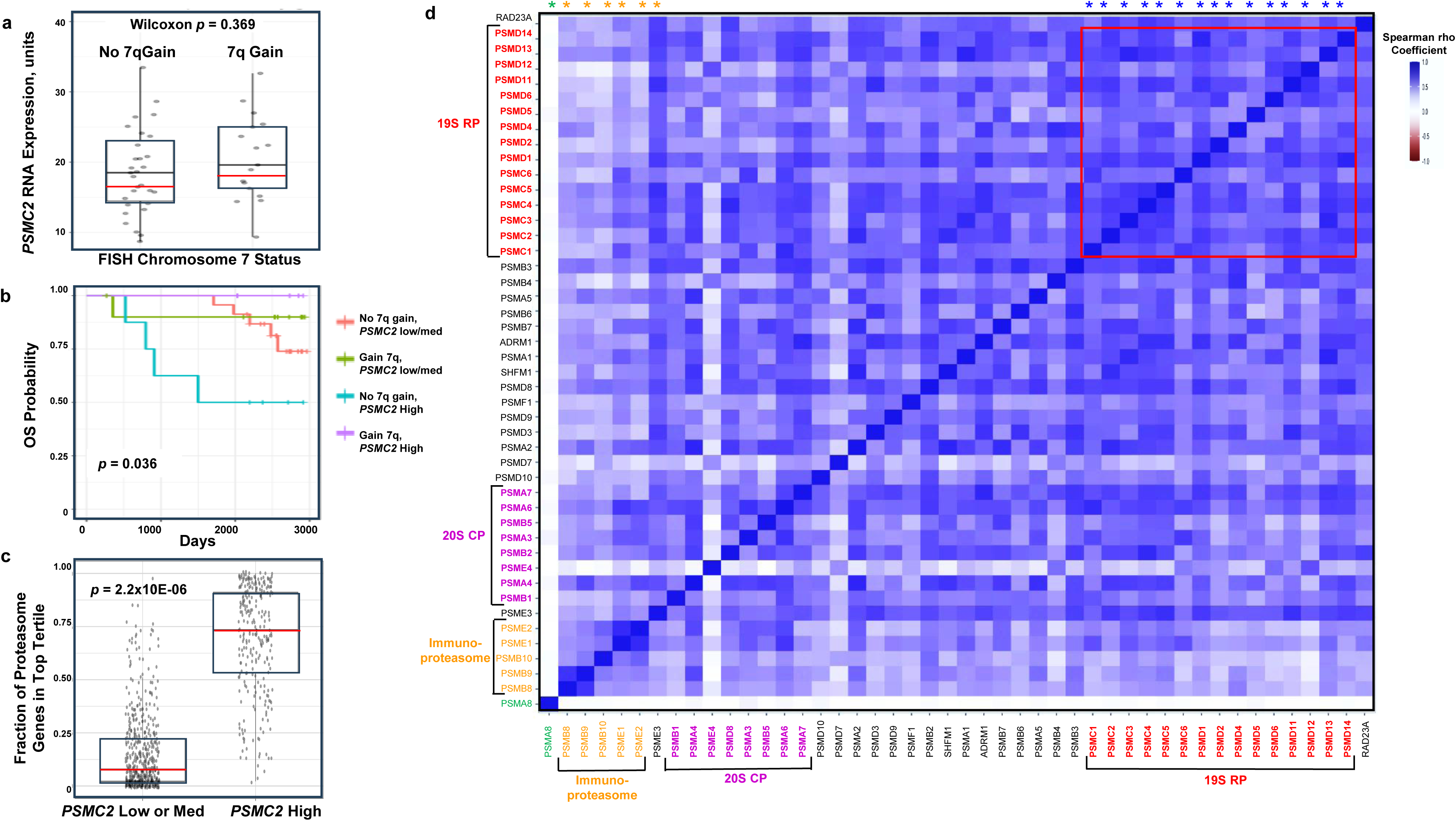
*PSMC2* is a unique, independent prognostic factor of Btz resistance. **a**. Lack of *PSMC2* expression correlation with chromosome 7q gain. Wilcoxon analysis compared *PSMC2* expression levels from CoMMpass dataset patients with gain 7q (n=17) and without gain of 7q (n=31). Results revealed no difference in *PSMC2* expression between these groups (p=0.369). **b.** K-M analysis comparing patients from the CoMMpass dataset without gain 7q with low/med *PSMC2* tertile of expression (n=23), patients without gain 7q with high *PSMC2* tertile expression (n=8), patients with gain 7q with low/med *PSMC2* tertile expression (n=11), and patients with gain 7q with high *PSMC2* tertile expression (n=6) revealed a statistically significant difference in OS by the log-rank test with p=0.036. **c. Fraction of proteasome genes expressed in the highest tertile** of relative RNAseq expression compared between *PSMC2* low/med and *PSMC2* high patients using a Wilcoxon rank sum test with continuity correction. Results revealed that the *PSMC2* high tertile patients were more likely to exhibit high tertile expression of additional proteasomes genes (p=2.2 x10^-6^). **d**. Co-expression of proteasome genes was assessed using a Spearman’s Rho analysis. *PSMA8* (green star) is expressed only in spermatoproteasomes and did not correlate with any other proteasome gene. Immunoproteasome-specific genes are denoted by gold stars, while proteasome 19S RP *PSMC* and *PSMD* genes are indicated in blue.

Prior work reported that the coordinated induction of UPS components is potentially responsible for enhanced protein degradation observed under stress conditions^31,32^. While higher *PSMC2* expression specifically correlated with reduced clinical response to Btz, it has been observed that certain proteasome genes may also function as an evolutionarily conserved network. We performed a co-occurrence analysis to test if increased *PSMC2* expression was coordinated with increased expression of other proteasome genes (Fig. 2c). High expression for each gene was defined as the top tertile within that given patient cohort, and the frequency of simultaneous high-expression states across genes then quantified. The analysis revealed that patients who harbored myeloma cells with *PSMC2* expression in the highest tertile were much more likely to also express higher levels of other proteasome genes, specifically *PSMC* and *PSMD* (Fig. 2c; p=2.2 x10^-6^). A Spearman rank correlation analysis revealed further concordance between the increase in *PSMC2* expression with the expression of individual *PSMC* and *PSMD* genes (Fig. 2d). The Spearman analysis revealed tightly coordinated transcriptional control of multiple *PSMC* and *PSMD* genes in MM patients. We focused downstream studies on *PSMC2* since it was the sole proteasome gene that correlated with clinical response to Btz, PFS, and OS using patient datasets from both trials. Hence, we posited that *PSMC2* overexpression may create unique synthetic lethal vulnerabilities that make it a prime target for precision therapies to overcome Btz resistance.

### *PSMC2-*overexpression increases 19S RP and 26S proteasome levels

PSMC2 plays a crucial role in the assembly of 19S RPs as well as its physical association with 20S CPs to form mature 26S proteasome complexes. PSMD5 (S5b) is an assembly chaperone that contacts both PSMC1 and PSMC2 and promotes their pairing within a PSMD5-PSMC2-PSMC1-PSMD2 module, but PSMD5 is displaced during maturation and is not incorporated into mature 26S proteasomes^18,33^. During assembly, the six PSMC ATPases are delivered as three chaperone-bound modules (PSMD5-PSMC1-PSMC2; PSMD9-PSMC6-PSMC3; and PSMD10-PSMC4-PSMC5)^34^. PSMD2 is retained in the mature base, where it makes extensive contacts with the outer face of the ATPase ring^33^, so the PSMC2-PSMC1-PSMD2 species detected here by co-immunoprecipitation represents the assembled, chaperone-free form. We expressed FLAG-tagged PSMC2 and PSMC6 in U266 and ARH-77 cells. FLAG-tagged PSMC2 and PSMC6 subunits co-immunoprecipitated with PSMC1 and PSMD2 as indicated by western blot (Fig. 3a). However, non-denaturing gel electrophoresis and western blotting indicated that FLAG-tagged PSMC2, but not FLAG-tagged PSMC6, was incorporated into 19S RPs (Fig. 3b). (PSMC6-FLAG alone was readily detected by western in lysates). The results further supported focusing studies on *PSMC2*, since overexpression of *PSMC2*, but not *PSMC6*, led to a significant increase in the level of 19S RPs in MM cell lines.

**Fig. 3.**
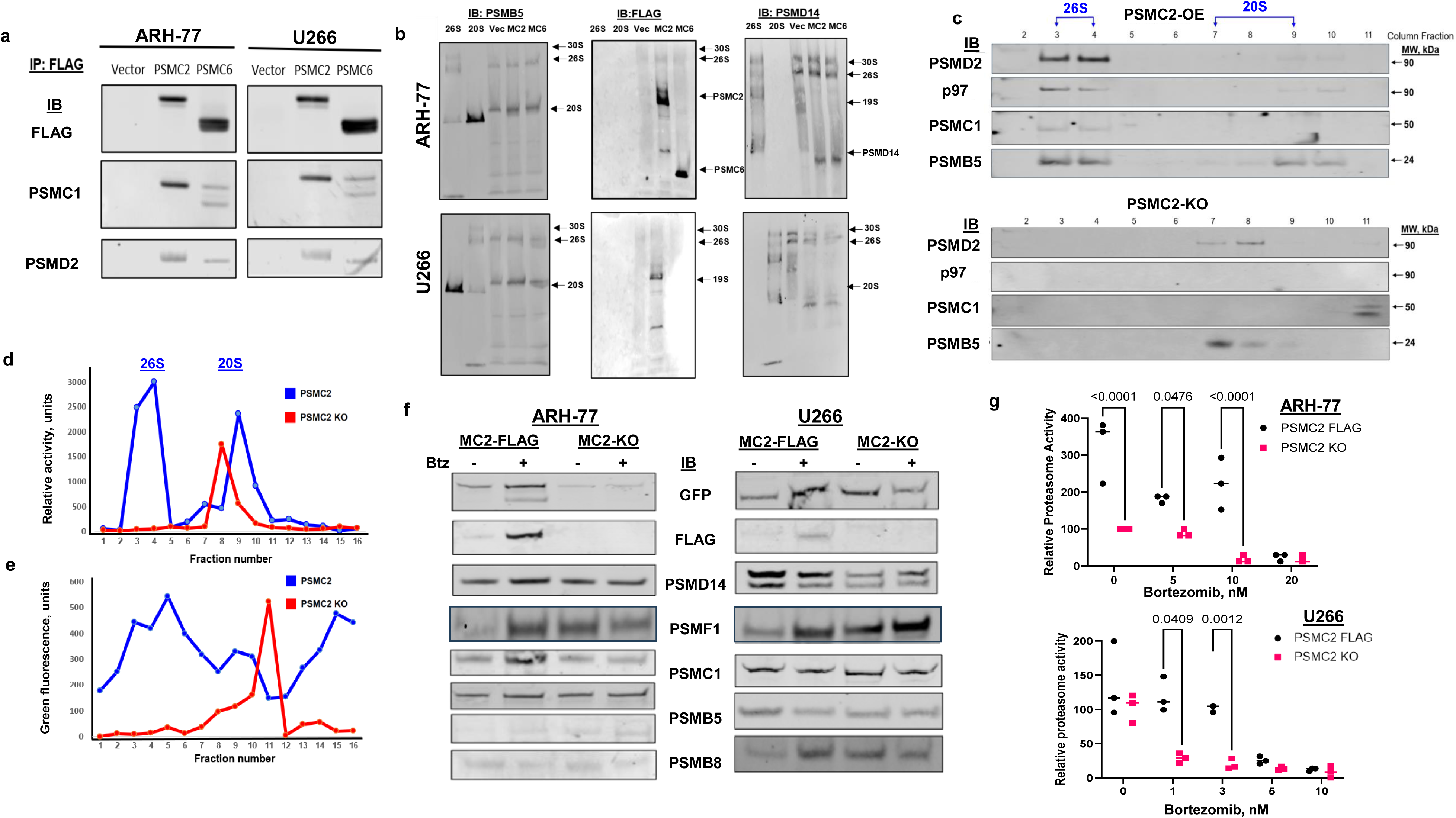
*PSMC2*-overexpression increases the levels of 19S RPs and 26S proteasomes. **a.** FLAG-tagged-PSMC2 was overexpressed in ARH-77 and U266 cells, cell lysates were prepared and proteasomes immunoprecipitated using a FLAG-sepharose. Western blots indicated that FLAG-tagged PSMC2 co-immunoprecipitated with PSMC1 and PSMD2. Immunoprecipitates of FLAG-tagged PSMC6 indicated a weaker association of PSMC6 with PSMC1 and PSMD2 **b**. Non-denaturing PAGE (3-8%) analysis of FLAG immunoprecipitates obtained from *PSMC2-FLAG* and PSMC6-FLAG expressing ARH-77 and U266 cells. Purified human 26S and 20S proteasomes were used as controls to monitor protein migration. As expected, PSMB5 was detected in both 26S and 20S proteasomes. PSMD14 is an essential structural and catalytic component of the 19S RP and the 26S proteasome, responsible for removing ubiquitin chains from protein substrates. Blots indicated that *PSMC2-FLAG* was incorporated into 19S and 26S complexes within both cell lines. Overexpressed FLAG-tagged PSMC6 was not incorporated into 19S and 26S proteasomes as readily as FLAG-tagged PSMC2. **c**. Twin-Strep-tagged-PSMD14 was used to immunoprecipitate 26S proteasomes from U266 cells that either expressed *PSMC2-FLAG* or had undergone *PSMC2*-*KO*. After TEV digest, the eluate was resolved by analytic SEC and column fractions were analyzed by western blotting. Greater levels of PSMC1, PSMC2, PSMD2, PSMB5 and p97 were detected in fractions 2-4 (Purified human 26S proteasomes eluted in fractions 2-4, while 20S proteasomes eluted in fractions 8-10). **d**. Proteasome hydrolysis of the fluorogenic peptide Suc-LLVY-AMC was measured in SEC column fractions. Activity was detected in fractions 2-4 from PSMC2-FLAG cells but not in fractions 2-4 from PSMC2-KO cells. **e.** *PSMC2*-overexpressing and *PSMC2-KO* U266 cells were engineered to express the PSMD14-TEV-affinity tag and transfected with a plasmid that expressed p97-GFP. Proteasomes were immunoprecipitated as previous and eluted proteins separated by SEC. Green fluorescence in each column fraction was then measured and revealed higher levels of GFP-p97 in fractions 2-4 in PSMC2-FLAG cells compared to *PSMC2*-*KO* cells. **f**. U266 and ARH-77 cells expressing affinity-tagged PSMD14, p97-GFP and either PSMC2-FLAG or *PSMC2*-*KO* were treated with 10 nM Btz for 18 h, lysed, proteasomes affinity-purified and analyzed by western blot to reveal increased GFP (p97-GFP) and PSMC1, and reduced PSMF1, after Btz challenge in *PSMC2-FLAG* cells relative to PSMC2-KO cells. **g.** Proteasomes from ARH-77 and U266 cells expressing PSMC2-FLAG or *PSMC2*-*KO* were affinity purified and treated with Btz. Proteasome activity was measured. In ARH-77 cells, baseline activity was higher in *PSMC2-KO* than in *PSMC2-FLAG* cells, whereas baseline activity was comparable between the two in U266 cells; with increasing Btz concentration, *PSMC2-FLAG* cells retained greater activity per dose in both cell lines. Differences in means were compared using two way ANOVA testing with Sidak’s multiple comparison testing. p-values are given.

Given that *PSMC2* plays a key role in the formation of 19S RPs, we hypothesized that increased *PSMC2* expression may also increase formation of 26S proteasomes. To interrogate this possibility, MM cells were transduced with lentiviral sequence that expressed the 26S proteasome non-ATPase 14 (*PSMD14*) gene fused to the cleavable tobacco etch virus (TEV) protease site followed by an affinity tag. (U266 cells expressed Twin-Strep-tagged-PSMD14 while ARH-77 cells expressed GST-tagged-PSMD14). Cells were also transfected to express p97-GFP. Lysates were prepared under non-denaturing conditions, incubated with affinity resin, bound protein released by TEV digestion, and proteins probed by western blotting. Lysates were analyzed by non-denaturing gel electrophoresis and probed with antibodies specific to 19S RP and 20S CP subunits (Fig. 3b). Results indicated that *PSMC2*-overexpressing cells contained a higher level of 19S RPs than pLKO (empty vector) control or *PSMC6*-overexpressing cells. The results revealed an unforeseen critical role of *PSMC2* on 19S RP formation, and hence, 26S proteasome assembly. Lysates from *PSMC2*-overexpressing and *PSMC2*-*KO* cells were applied to affinity-resin, released by TEV digestion, and purified by analytic SEC. Activity assays demonstrated that purified 26S proteasomes eluted in fractions 2-4, while 20S proteasomes eluted in fractions 8-10. Western blotting indicated greater levels of PSMC1, PSMC2, PSMD2, PSMB5, and p97 in fractions 2-4 from PSMC2-overexpressing cells relative to the same fractions from *PSMC2-KO* cells (Fig. 3c; Fig. S11). Proteasome activity was also greater in fractions 2-4 of *PSMC2-*overexpressing cells relative to *PSMC2-KO* cells (Fig. 3d). We also detected increased green fluorescence, a marker of p97-GFP, in fractions 2-4 of *PSMC2-* overexpressing cells to support physical association of p97 with 26S proteasomes (Fig. 3e).

PSMF1/hPI31 is a highly conserved protein ubiquitously expressed across human tissues that has been reported to inhibit 20S proteasome peptide hydrolysis *in vitro* through its C-terminal proline-rich domain^35–37^. PSMF1 has been proposed to physically block or compete with 19S RPs and PA28α/b complexes for docking onto the α-rings of the 20S CP. However, PSMF1 has also been reported to stimulate 26S proteasome-mediated proteolysis *in vitro*^38^. 26S proteasome complexes affinity-purified from *PSMC2*-overexpressing cells demonstrated significantly higher amounts of p97, while the level of PSMF1 bound to these proteasomes was low. (Fig. 3f). Conversely, the association of p97 with proteasomes from *PSMC2-KO* cells was low, while the association of PSMF1 was higher relative to *PSMC2*-overexpressing cells. The activity of proteasomes isolated from *PSMC2*-overexpressing cells was not only greater at baseline, but also following Btz treatment compared to the proteasomes isolated from *PSMC2*-*KO* cells (Fig. 3g). It is plausible that PSMC2 contributes to a context-dependent association of p97 and PSMF1 with proteasomes to fine-tune proteolytic activity during cellular stress^39,40^.

### *PSMC2-KO* promotes Ub-chain accumulation on proteasomes after Btz treatment

We hypothesized that proteasomes isolated from *PSMC2-*overexpressing cells could process ubiquitinated substrates more efficiently than proteasomes from *PSMC2-KO* cells following Btz treatment. U266 and ARH-77 cells were treated with Btz, incubated with the crosslinker DSP, proteasomes affinity-purified^41^, and probed for covalently attached linkage-specific Ub chains^42^. Western blotting revealed that the relative level of the canonical degradation-associated K48 and K63-specific Ub chains associated with proteasomes purified from *PSMC2-KO* cells was much greater than that associated with proteasomes purified from *PSMC2*-overexpressing cells following Btz treatment (Fig. 4; Fig. S12). While proteasomes lacking *PSMC2* recognize and dock ubiquitinated substrates, the deubiquitination, unfolding, translocation, and cleavage of these substrates appears impaired and leads to less efficient processing. The direct recruitment of p97 to 19S RPs has been proposed to provide additional upstream mechanical energy to relieve proteasomal stalling^43^. Our results demonstrate that proteasomes from *PSMC2-KO* cells not only lack p97 but accumulate ubiquitinated substrates upon Btz challenge. In contrast, proteasomes from *PSMC2*-overexpressing cells exhibit greater catalytic activity and do not accumulate ubiquitinated substrates.

**Fig. 4.**
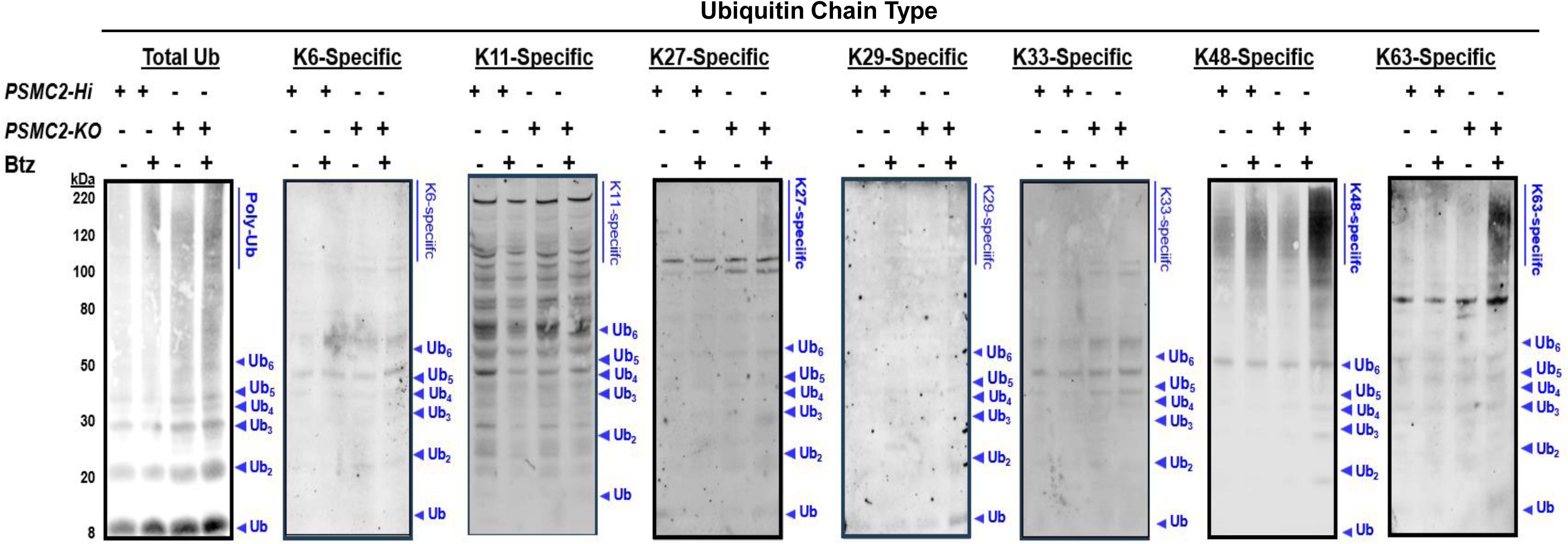
Btz promotes Ub chain accumulation on proteasomes in *PSMC2-KO* cells. *PSMC2*-overexpressing and *PSMC2-KO* U266 cells were treated with 10 nM Btz for 18 h, crosslinked with DSP, lysed in non-denaturing buffer and affinity purified using the PSMD14-TEV-GST system. Proteins were boiled in sample buffer, separated on 4-12% NuPAGE SDS gels and probed by western blot using antibodies specific to each homotypic Ub chain type; the Ub chain linkage is indicated above each panel. Similar results were observed using ARH-77 lysates (Fig. S12).

### *PSMC2*-overexpression is synthetic lethal with p97 ATPase inhibition

Synthetic lethal interactions represent a tactical means to therapeutically exploit differences between cancer and healthy cells^44^. Since MM cells are exquisitely sensitive to agents that disrupt proteostasis, we queried the effect of ER-targeting pharmacologics on *PSMC2*-overexpressing cells^45^. We utilized U266 cells which express relatively high levels of *PSMC2* compared to other MMCLs (Fig. S13). Cells were transfected with vectors that expressed either *pLKO* (control) or *PSMC2*-*FLAG* (Fig. 5a). We identified 4 compounds that selectively reduced the viability of *PSMC2*-overexpressing cells >2-fold greater than that of controls. These compounds were: p97/VCP inhibitor CB-5339, eIF2α integrated stress response inhibitor ISRIB, IRE1α inhibitor KIRA6, and Hsp90 inhibitor PF-04929113 (Fig. 5b). p97 is a multifunctional, AAA^+^-type ATPase involved in protein quality control and has been linked to the UPS^46,47^. A prior study identified a compound structurally similar to DBeQ that exerted anti-myeloma activity, in part through p97/VCP inhibition and suggested that the compound induces ER stress^48^. DBeQ inhibits p97 to block Ub-fusion and ER-associated protein degradation as well as autophagosome maturation. Dual inhibition of p97 and the proteasome also terminally disrupted ER configuration and induced apoptosis in MMCLs^49^.

**Fig. 5.**
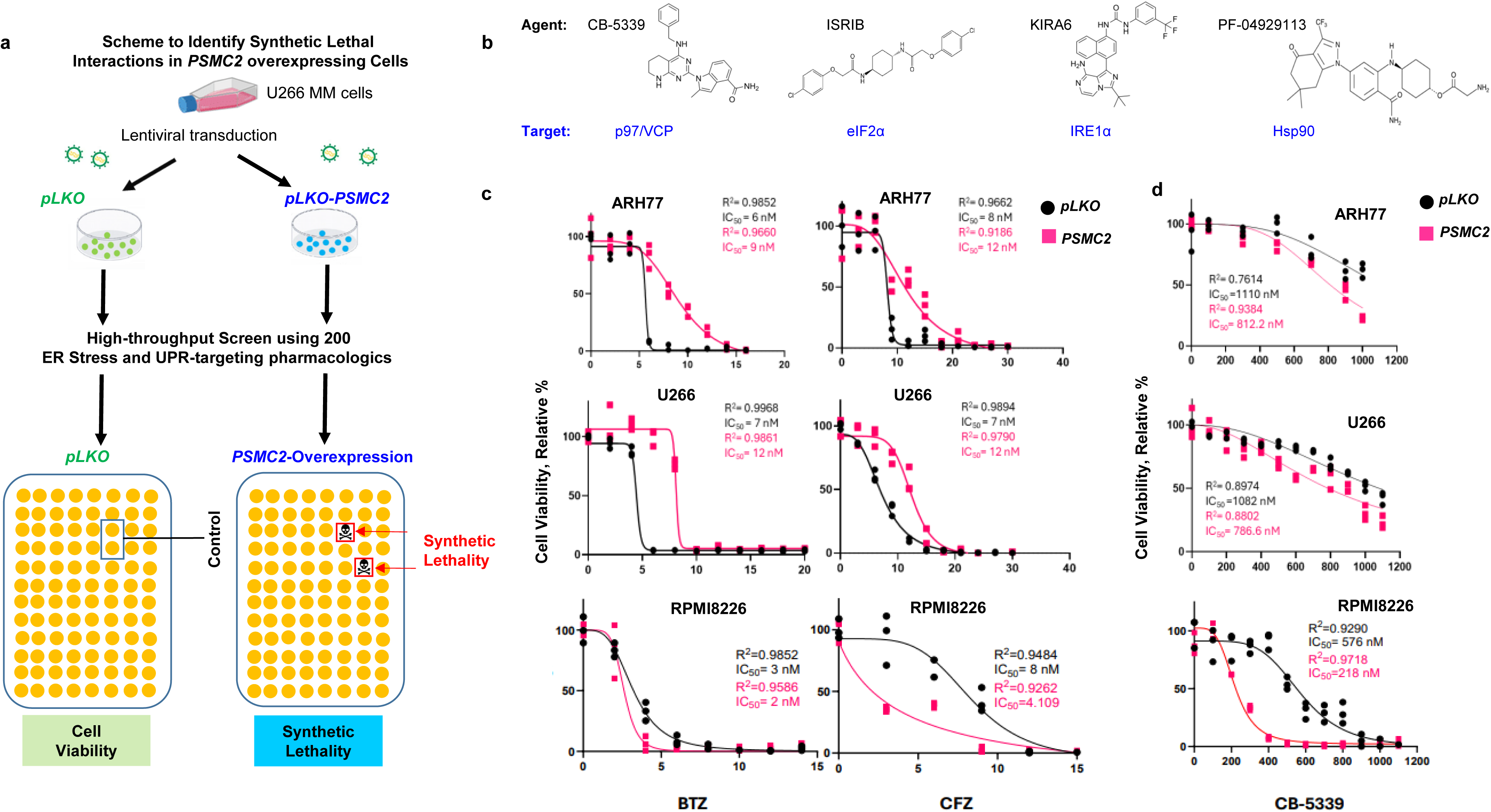
*PSMC2*-overexpression is synthetic lethal with the p97 ATPase inhibitor CB-5339. **a**. Scheme for high throughput pharmacologic screens to detect synthetic lethal interactions in *PSMC2*-overexpressing cells compared to myeloma cells expressing the empty vector control *pLKO*. U266 cells were transduced with either *pLKO*- or *PSMC2-FLAG* constructs and subsequently challenged with a library of 200 small molecules that target effectors of the ER stress and unfolded protein response (UPR). Molecules were added to cells plated in 96 well plates at concentrations of 0.3, 1 and 3 uM. After three days of treatment, relative cell viability was assayed using the XTT assay. Hits were identified as compounds that increased cell killing in *PSMC2*-overexpressing cells 2-fold greater than that measured in *pLKO*-expressing cells. **b**. Chemical structures of the top four HTS hits identified: CB-5339 (p97 inhibitor), ISRIB (EIF2α inhibitor), KIRA6 (IRE1α inhibitor) and PF-04929113 (HSP90 inhibitor). **c.** pLKO-(black) or PSMC2-transduced (pink) cells, as annotated above each panel, were treated with increasing concentrations of Btz or Cfz, for 3 days and viability was measured using the XTT assay. Results are expressed as relative absorbance compared to untreated cells with n=3 in all cases. Data was analyzed using nonlinear regression analysis in Graphpad. R2 and LD50 values are reported. **d.** *pLKO* and *PSMC2*-transduced MM cells were treated with CB-5339 for three days before viability was measured using the XTT assay. Results are expressed as relative absorbance compared to untreated cells with n=3 in all cases. Data was analyzed using nonlinear regression analysis in Graphpad. R2 and LD_50_ values are reported.

CB-5339 is a selective, p97/VCP ATPase inhibitor which was prioritized based upon direct linkage to the UPS, on-target efficacy, oral availability, and promising early phase clinic trial results^50–52^. We validated the findings in RPMI-8226 cells that express lower *PSMC2 per* the Cancer Cell Line Encyclopedia, as well as ARH-77 cells. Interestingly, PSMC2-overexpressing ARH-77 and U266 cells demonstrated higher LD_50_ values for PIs than pLKO controls, whereas this relationship was reversed in RPMI-8226 cells; PSMC2-overexpressing cells exhibited increased CB-5339 sensitivity (Fig. 5c, d).

### *p97* KD selectively sensitizes *PSMC2*-overexpressing cells

Since *PSMC2*-overexpressing cells were highly sensitive to pharmacologic inhibition of p97 ATPase activity, we determined whether *PSMC2*-overexpressing cells were also sensitive to *p97* genetic ablation. *p97*-*KO* is deleterious at the cellular level which required construction of a *Tet*-inducible conditional knockdown (KD) system to model the effect of *p97* loss on cellular sensitivity to CB-5339. Tet-inducible p97-KD cell lines were engineered in parental MMCLs that expressed either high *PSMC2* (U266) or low basal *PSMC2* (RPMI-8226) cells (Fig. 6a). Following *Tet*-stimulation to KD *p97* expression, marked cell death of *PSMC2*-overexpressing cells was detected relative to untreated controls (Fig. 6b). Following *Tet* stimulation and *p97* KD, *PSMC2*-overexpressing cells were more sensitive to CB-5339 and Btz treatment relative to pLKO-transfected controls (Fig. 6c, d).

**Fig. 6.**
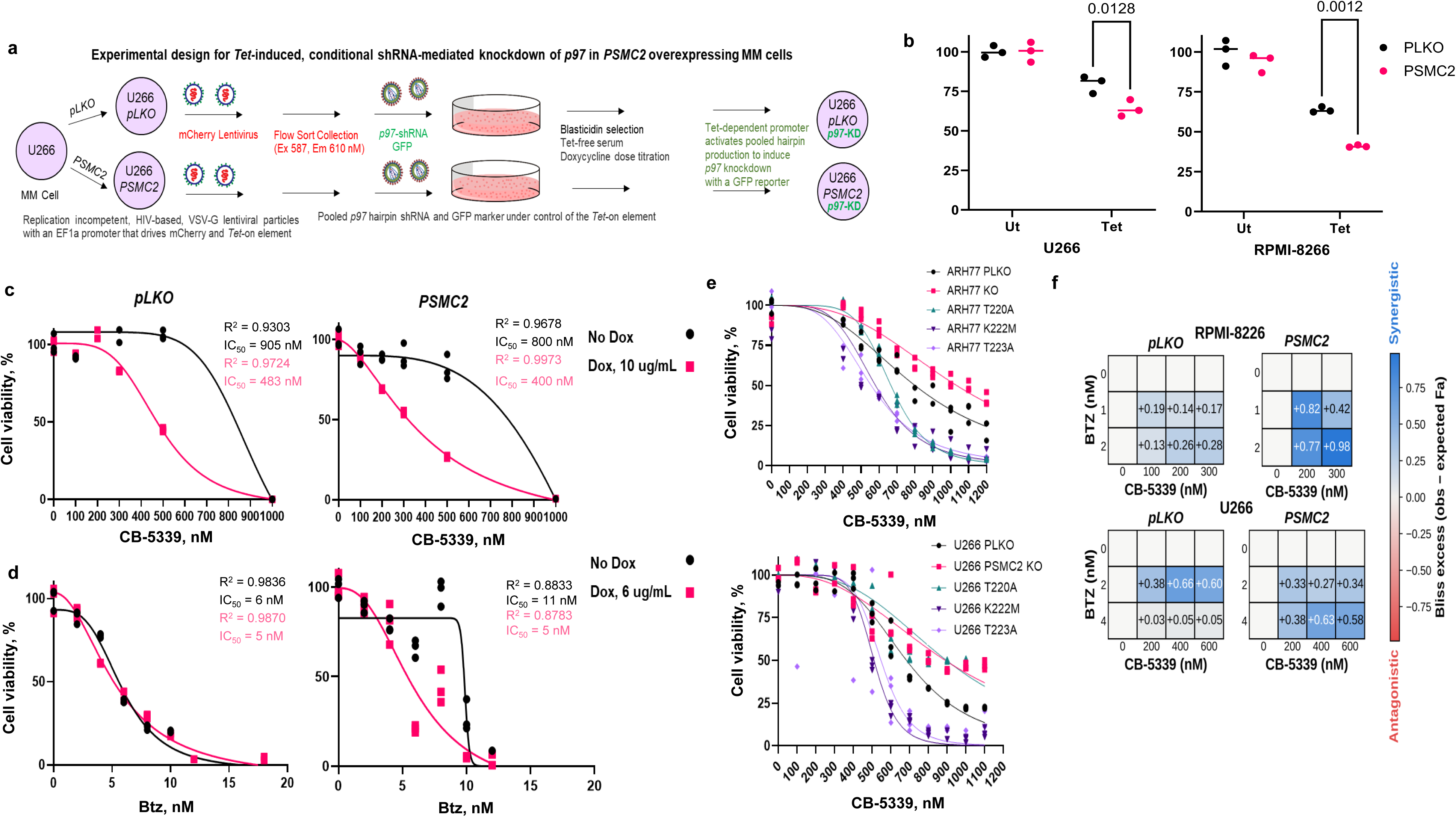
*PSMC2*-overexpressing cells are more sensitive to p97 KD and Btz or CB-5339 treatment. **a.** Schema of conditional p97 KD. *pLKO* or *PSMC2-FLAG* expressing U266 cells were sequentially transduced to stably express a Dox-responsive transcription cofactor and selected by mCherry flow sorting. mCherry positive cells were transduced with 3 shRNAs to knockdown p97 and expressed GFP under control of a third-generation, blasticidin-selectable Tet-On promoter. Lines are designated as MMCL-*pLKO*-Cherry and MMCL-*PSMC2*-Cherry. **b**. U266 and RPMI vector and PSMC2 cherry cells were treated with 10 μg/ml Dox for 3 days and cell viability determined using the XTT assay. Viability of *pLKO* cells decreased by ∼30%, while viability of *PSMC2*-expressing cells decreased ∼50% after Dox treatment. Differences between Dox and no Dox treatment mean values were compared by Sidak’s multiple comparison testing (n = 3, *p*-values given). **c.** U266-pLKO-Cherry and U266-*PSMC2-Cherry* cells were pretreated with 10 μg/ml Dox for 3 days, at which point an equal number of cells were treated with increasing doses of CB-5339 for 3 days in triplicate prior to measuring cell viability using the XTT assay. A nonlinear curve was extrapolated with R2 and LD50 values reported. **d**. U266-*pLKO*-Cherry and U266-*PSMC2*-Cherry cells were pretreated with 6 μg/ml Dox for 3 days, at which point an equal number of cells were removed and treated with increasing doses of Btz for 3 days in triplicate prior to measuring cell viability using the XTT assay. A nonlinear curve was extrapolated with R2 and LD50 values reported. **e**. PSMC2-KO cells engineered to express the PSMC2 ATPase active-site point mutants T220A, K222M or T223A, alongside PSMC2-pLKO and PSMC2-KO controls, were treated with increasing doses of CB-5339 for 3 days prior to measuring cell viability using the XTT assay. **f**. RPMI-8226 and U266 *pLKO* and *PSMC2*-expressing cell lines were treated with indicated doses of Btz and CB-5339 and assayed for viability using the Alamar Blue assay. Bliss synergy was calculated where positive values reveal synergistic killing (n = 3 for each condition).

### PSMC2-mediated p97 inhibition sensitization is not dependent on ATPase activity

To test if *PSMC2*-mediated sensitization of MMCLs was dependent on PSMC2 ATPase activity, point mutations were generated at T220A, K222M, and T223A to disrupt the ATPase active site (Uniprot P35998). *PSMC2*-*KO* cells were then engineered to express each *PSMC2* point mutant and the effect of CB-5339 on cell viability determined (Fig. 6e). MM cells that expressed the *PSMC2* K222M and T223A mutants were more sensitive to CB-5339 treatment than *PSMC2*-pLKO and *PSMC2-KO* cells. The exquisite sensitivity to CB-5339 in cells that express *PSMC2* mutants suggests that these cells maintain p97-dependency that is further impacted by defective PSMC2-ATPase subunits.

### Btz and CB-5339 synergize better in *PSMC2*-overexpressing cells

Cells were treated with Btz and CB-5339 alone and pair-wise combinations across a dose matrix and cell viability measured. The analysis revealed Bliss excess in *PSMC2*-expressing cells relative to pLKO-expressing cells with the drug combination in all conditions, consistent with prior results^49^ (Fig. 6f).

### *In vivo* synergistic activity of CB-5339 and Btz against myeloma xenografts

To address the *in vivo* anti-myeloma effect of CB-5339, ARH-77 cells that expressed either *pLKO* or *PSMC2* were xenografted into NSG mice (Fig. 7a; Fig. S14). Following formation of palpable tumors, mice were treated with either vehicle, CB-5339, Btz or both. Co-treatment reduced bioluminescence of xenografts in mice that had received *PSMC2*-overexpressing cells compared to vehicle and CB-5339 alone, but not Btz alone (Fig. 7b). However, CB-5339 and Btz co-treatment preferentially prolonged the survival of mice injected with *PSMC2-*overexpressing cells compared to the other treatments (Fig. 7c, d). Weight loss, neurologic, musculoskeletal, and hematopoietic adverse effects were not observed.

**Fig. 7.**
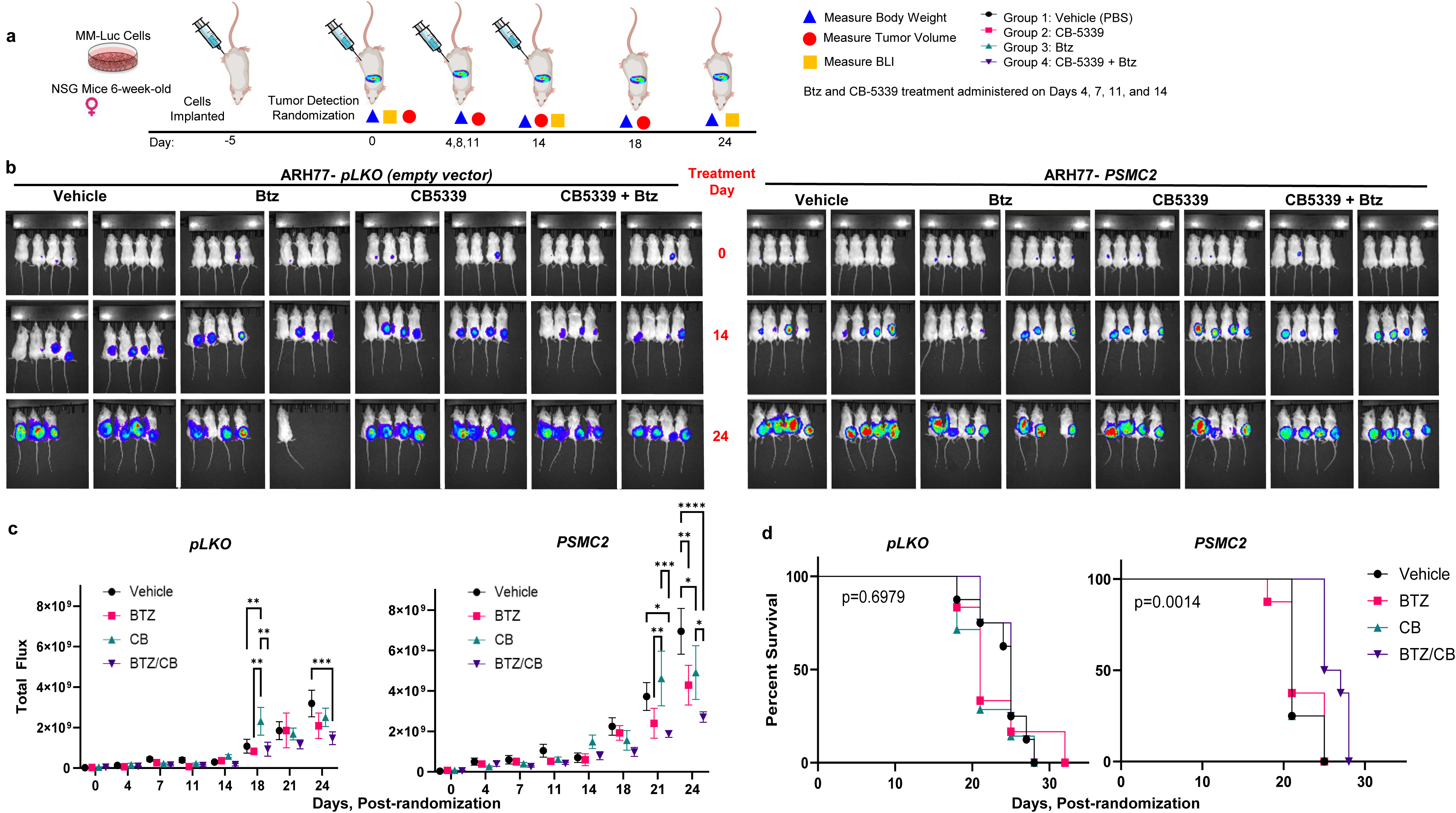
*PSMC2*-overexpression sensitizes MM cells to combined CB-5339 and Btz therapy *in vivo*. **a**. Experimental schema: 64 female NSG mice > 6 weeks of age were injected with 1.5x10^6^ ARH77-pLKO-LUC or ARH77-PSMC2-LUC cells subcutaneously (sc). Five days after injection, mice were injected with luciferin sc. Ten minutes later, total photon flux was quantified using a Spectrum IVIS with a whole mouse gate using LivingImage software. For each cell line, mice were randomized into four treatment groups of 8 mice (eight groups, 64 mice total). [Two mice in the *pLKO* Btz group and one in the CB-5339 group did not develop tumors and were excluded from analysis (n=6 and n=7 for these groups, respectively)]. Groups were treated with 0.5 mg/kg Btz sc, 45 mg/kg CB-5339 by oral gavage or combination therapy twice weekly for 2 weeks. Mice were monitored twice weekly for tumor size by BLI and for morbidity. **b.** Normalized total photon flux was analyzed at the start of therapy (t=0), and at day 14 and day 24. **c**. Average photon flux/treatment group plotted over time. Dunnett’s multi-comparison analysis compared total flux between treatment groups with exact p-values given in the figure. n=8 for each group except n=6 for pLKO Btz and n=7 for pLKO-CB-5339. **d**. K-M survival analysis of either ARH77-*pLKO-LUC* or ARH77-*PSMC2-LUC* injected mice treated with vehicle, 0.5 mg/kg Btz twice weekly for 2 weeks, 45 mg/kg CB-5339 twice weekly for 2 weeks or combination treatment. Median OS improved in ARH77-*PSMC2-LUC* mice from 21 days in vehicle to 24 days with combination treatment. OS was analyzed by log-rank test, revealing a statistically significant difference between ARH77-*PSMC2-LUC* treatments (p = 0.0014). Mice were euthanized with CO_2_ inhalation to minimize distress at the earliest sign of predefined endpoints to prevent suffering.

### Synergy of CB-5339 with Btz in patient samples

Chou-Talalay analysis indicated that CB-5339 and Btz co-treatment of CD138^+^ cells from patients previously exposed to Btz generated synergistic cytotoxicity^53^ (Fig. S15a; Table S4). Co-treatment of MMCLs with CB-5339 and Btz also synergistically targeted MM cells and overcame Btz-resistance (Fig. S15b). Btz and CB-5339 combination treatment increased caspase-3 cleavage in MMCLs, consistent with an apoptotic effect (Fig. S16a, b), but did not appreciably reduce PBMC viability.

## Discussion

Our study has defined a synthetic lethal vulnerability associated with high expression of the proteasome ATPase, *PSMC2*. MM cells harbor an extraordinarily complex genomic landscape that bestows phenotypic diversity, unpredictable treatment response, and heterogeneous clinical outcomes. *PSMC2* may serve as practical and biologically interpretable single-gene proxy for a broader adverse proteasome-expression state, rather than a statistically superior replacement for a multi-gene signature. The results support a more nuanced and defensible argument in that *PSMC2* is a robust adverse prognostic marker, ranks highest among the proteasome genes, particularly for OS, and captures much of the prognostic information contained in much larger proteasome signatures. Its principal advantage is not superior statistical performance compared to multigene signatures, but rather parsimony, biological relevance, and potential ease of clinical translation.

p97 functions as a molecular motor to convert energy from ATP hydrolysis into physical energy to unfold and thread substrates through the hexameric channel and attenuate bottlenecks in substrate processing^54–56^. While both p97 and proteasomes are abundant, only a fraction of proteasomes are processing substrates at a given moment and the steady-state population of p97 hexamers bound to proteasomes appears minute. Our results suggest that under homeostatic conditions, p97 is associated with a subset of proteasomes in a transient, hybrid configuration that is unable to withstand conventional purification methods. *PSMC2-*overexpression leads to either a greater proportion of p97 complexed with proteasomes or extends the length of time during which p97 associates with proteasomes. CB-5083 is another p97 inhibitor reported to decrease viability in MMCLs and patient-derived MM cells, including those with background PI-resistance^57^. CB-5083 may block p97-dependent retro-translocation of the transcription factor, Nrf1, which transcribes proteasome subunit genes following PI exposure. Cryo-EM and mass spectroscopy studies should precisely define how physical association of p97 with the 19S RP provides additional mechanical energy to increase substrate unfolding and entry into the inner chamber of the 20S CP to benefit the processing of ubiquitinated clients. Other non-canonical proteasome structural adaptations have been reported to enhance proteasome-dependent proteolysis^58–61^. The discovery of unforeseen, clinically-relevant structural adaptations that expand the functional repertoire of proteasomes provides insights to further improve cancer treatment.

## Acknowledgements.

Funding - Research was supported by NIH R01 (5R01AI139141), the UH Seidman Cancer Center, the Vinney Foundation and the Case Comprehensive Cancer Center (CCC). This research was supported by the Translational Research Shared Resources of the Case CCC (P30 CA#43703). The authors state that the Life Technologies Attune NxT Flow Cytometer was acquired through an NIH Shared Instrumentation Grant and, as such, we are required by NIH Regulations to cite the assistance of the Case CCC Cytometry & Microscopy Shared Resource and NIH Grant S10-NIH OD021559.

## Data availability

Gene expression and clinical data from the APEX trial are available from the NCBI Gene Expression Omnibus under accession GSE9782. Data from the Multiple Myeloma Research Foundation CoMMpass study (release IA22) are available to qualified investigators through the MMRF Researcher Gateway (https://research.themmrf.org). Gene dependency, CRISPR/CAS9 knockout, RNAi and copy number data from Project Achilles are available through the DepMap portal (https://depmap.org).

## Declaration of competing interests

The authors declare no competing interests.

## Contributions

JJD, EM, JIH and ERC developed the concept. JIH, ERC, JJD, WH, MS, and JDC performed experiments. JJD, JIH, EM, ERC, WH, DJT, NM, MS, NCM, KCA and JDC analyzed the data. JJD wrote the initial version of the manuscript. JJD, ERC, EM, MS and JIH prepared the figures. All remaining authors edited subsequent versions, made substantial contributions and approved the final version.

## Ethics approval

The study was approved by the Institutional Animal Care and Utilization Committee at Case Western Reserve University.

## Notes

### Competing Interest Statement

The authors have declared no competing interest.

## References

1. Malard F, Neri P, Bahlis NJ, et al. Multiple myeloma. Nat Rev Dis Primers 2024; 10, 45. doi10.1038/s41572-024-00529-7.

2. Samur MK, Aktas Samur A, Fulciniti M, et al. Genome-Wide Somatic Alterations in Multiple Myeloma Reveal a Superior Outcome Group. J Clin Oncol. 2020;38(27):3107–18. doi: 10.1200/JCO.20.00461. PMID: 32687451; PMCID: PMC7499613.

3. Bolli N, Avet-Loiseau H, Wedge D, et al. Heterogeneity of genomic evolution and mutational profiles in multiple myeloma. Nat Commun. 5, 2997 (2014). doi10.1038/ncomms3997.

4. Maura F, Boyle EM, Coffey D, et al., Genomic and immune signatures predict clinical outcome in newly diagnosed multiple myeloma treated with immunotherapy regimens. Nat Cancer. 2023;4(12):1660–74. doi: 10.1038/s43018-023-00657-1.

5. Skerget S, Penaherrera D, Chari A. et al. Comprehensive molecular profiling of multiple myeloma identifies refined copy number and expression subtypes. Nat Genet. 2024; 56, 1878–89. 10.1038/s41588-024-01853-0.

6. Cowan AJ, Green DJ, Kwok M, et al. Diagnosis and management of multiple myeloma: A Review. JAMA. 2022;327(5):464–77. doi: 10.1001/jama.2022.0003. PMID: 35103762.

7. Rajkumar SV. Multiple myeloma: 2022 update on diagnosis, risk stratification, and management. Am J Hematol. 2022;97(8):1086–107. doi: 10.1002/ajh.26590. PMID: 35560063; PMCID: PMC9387011.

8. Mateos MV, Ludwig H, Bazarbachi A, et al., Insights on multiple myeloma treatment strategies. Hemasphere. 2018;3(1):e163.

9. Malek E, Rana, PS, Swamydas M, et al. The TGFβ type I receptor kinase inhibitor vactosertib in combination with pomalidomide in relapsed/refractory multiple myeloma: a phase 1b trial. Nat. Commun. 2024, 15(1), 7388, doi: 10.1038/s41467-024-51442-2.

10. Lonial S, Popat R, Hulin C, et al., Iberdomide plus dexamethasone in heavily pretreated late-line relapsed or refractory multiple myeloma (CC-220-MM-001): a multicentre, multicohort, open-label, phase 1/2 trial. The Lancet Haematology. 2022;9(11):e822–32.

11. Meermeier EW, Bergsagel PL, Chesi M. Next-Generation therapies for multiple myeloma. Annu Rev Cancer Biol. 2024;8:351–371. doi: 10.1146/annurev-cancerbio-061421-014236. PMID: 39364307; PMCID: PMC11449476.

12. Bo Kim K. Proteasomal adaptations to FDA-approved proteasome inhibitors: a potential mechanism for drug resistance? Cancer Drug Resist. 2021;4(3):634–45. doi: 10.20517/cdr.2021.27. PMID: 34308274; PMCID: PMC8297691.

13. Li F, Liu J, Fu Y. Acquired Bortezomib Resistance in Multiple Myeloma: From Mechanisms to Strategy. Curr Treat Options Oncol. 2024;25(11):1354–1365. doi: 10.1007/s11864-024-01273-6. PMID: 39432172.

14. Besse A, Besse L, Kraus M, et al. Proteasome Inhibition in Multiple Myeloma: Head-to-Head Comparison of Currently Available Proteasome Inhibitors. Cell Chem Biol. 2019;26(3):340–351.e3. doi: 10.1016/j.chembiol.2018.11.007. PMID: 30612952.

15. Finley D. Recognition and processing of ubiquitin-protein conjugates by the proteasome. Annu Rev Biochem. 2009;78:477–513. doi: 10.1146/annurev.biochem.

16. Arkinson C, Dong KC, Gee CL. et al. Mechanisms and regulation of substrate degradation by the 26S proteasome. Nat Rev Mol Cell Biol 2025; 26:104–22.

17. Martinez-Fonts K, Davis C, Tomita T, et al. The proteasome 19S cap and its ubiquitin receptors provide a versatile recognition platform for substrates. Nat Commun. 2020;11(1):477. PMID: 31980598; PMCID: PMC6981147.

18. Le Tallec B, Barrault MB, Guérois R, Carré T, Peyroche A. Hsm3/S5b participates in the assembly pathway of the 19S regulatory particle of the proteasome. Mol Cell. 2009;33(3):389–99. doi: 10.1016/j.molcel.2009.01.010. PMID: 19217412.

19. Liu Y, Chen H, Li X, Zhang F, et al. PSMC2 Regulates Cell Cycle Progression Through the p21/Cyclin D1 Pathway and Predicts a Poor Prognosis in Human Hepatocellular Carcinoma. Front Oncol. 2021;11:607021. PMID: 33718159; PMCID: PMC7952995.

20. Wang Y, Zhu M, Li J. et al. Overexpression of PSMC2 promotes the tumorigenesis and development of human breast cancer via regulating plasminogen activator urokinase (PLAU). Cell Death Dis 2021; 12:690.

21. Gillette TG, Kumar B, Thompson D, Slaughter CA, DeMartino GN. Differential roles of the COOH termini of AAA subunits of PA700 (19 S regulator) in asymmetric assembly and activation of the 26S proteasome. Journal of Biological Chemistry. 2008;283(46):31813–22.

22. Tiedemann RE, Zhu YX, Schmidt J, Shi CX, Sereduk C, Yin H. Identification of molecular vulnerabilities in human multiple myeloma cells by RNA interference lethality screening of the druggable genome. Cancer Res. 2012;72(3):757–68. doi: 10.1158/0008-5472. PMID: 22147262; PMCID: PMC3622723.

23. Nijhawan D, Zack TI, Ren Y, et al., Cancer vulnerabilities unveiled by genomic loss. Cell. 2012;150(4):842–54. doi: 10.1016/j.cell.2012.07.023. PMID: 22901813; PMCID: PMC3429351.

24. Shi CX, Kortüm KM, Zhu YX, et al. CRISPR genome-wide screening identifies dependence on the proteasome subunit PSMC6 for bortezomib sensitivity in multiple myeloma. Mol Cancer Ther. 2017;16: 2862–70. PMID:28958990; PMCID: PMC5796678.

25. Kröll-Hermi A, Ebstein F, Stoetzel C, Geoffroy V, Schaefer E, Scheidecker S. Proteasome subunit PSMC3 variants cause neurosensory syndrome combining deafness and cataract due to proteotoxic stress. EMBO Mol Med. 2020;12(7):e11861. doi: 10.15252/emmm.201911861. PMID: 32500975; PMCID: PMC7338805.

26. Haertle L, Hernandez HNC, Buenache NS, Simicek M, Snaurova R, Rapado I. Genetic alterations in members of the proteasome 26S subunit, AAA-Atpase (*PSMC*) gene family transmit proteasome inhibitor resistance in multiple myeloma by impacting the ADP/ATP binding pocket. Blood 2022; 140 (Supplement 1): 4318–9.

27. Bliss CI. The toxicity of poisons applied jointly. Ann Appl Biol. 1939;26(3):585–615. doi: 10.1111/j.1744-7348.1939.tb06990.x.

28. Tsherniak A, Vazquez F, Montgomery PG, et al. Defining a Cancer Dependency Map. Cell. 2017;170(3):564–76.e16. 10.1016/j.cell.2017.06.010.

29. Dempster JM, Boyle I, Vazquez F, et al. Chronos: a cell population dynamics model of CRISPR experiments that improves inference of gene fitness effects. Genome Biol. 2021;22(1):343. doi:10.1186/s13059-021-02540-7.

30. Zhou Y, Zhang Q, Stephens O, et al. Prediction of cytogenetic abnormalities with gene expression profiles. Blood. 2012;119(21):e148–50.

31. Gilda JE, Nahar A, Kasiviswanathan D, et al. Proteasome gene expression is controlled by coordinated functions of multiple transcription factors. J Cell Biol. 2024;223(8):e202402046. doi: 10.1083/jcb.202402046. PMID: 38767572; PMCID: PMC11104393.

32. Rousseau A, Bertolotti A. Regulation of proteasome assembly and activity in health and disease. Nat Rev Mol Cell Biol 2018; 19:697–712 doi10.1038/s41580-018-0040-z

33. Tomko RJ Jr, Hochstrasser M. Molecular architecture and assembly of the eukaryotic proteasome. Annu Rev Biochem. 2013;82:415–45. doi: 10.1146/annurev-biochem-060410-150257. PMID: 23495936.

34. Tomko RJ Jr, Hochstrasser M. Order of the proteasomal ATPases and eukaryotic proteasome assembly. Cell Biochem Biophys. 2011;60(1-2):13–20. doi: 10.1007/s12013-011-9178-4. PMID: 21461838; PMCID: PMC3256250.

35. McCutchen-Maloney SL, Matsuda K, Shimbara N, et al. cDNA cloning, expression, and functional characterization of PI31, a proline-rich inhibitor of the proteasome. Journal of Biological Chemistry. 2000 Jun 16;275(24):18557–65.

36. Uhlén M, Fagerberg L, Hallström BM, et al. Tissue-based map of the human proteome. Science. 2015;347(6220):1260419.

37. Zaiss DM, Standera S, Holzhütter H, Kloetzel PM, Sijts AJ. The proteasome inhibitor PI31 competes with PA28 for binding to 20S proteasomes. FEBS letters. 1999;457(3):333–8.

38. Bader M, Benjamin S, Wapinski OL, et al. A conserved F box regulatory complex controls proteasome activity in Drosophila. Cell. 2011;145(3):371–82.

39. Clemen CS, Marko M, Strucksberg KH, et al. VCP and PSMF1: Antagonistic regulators of proteasome activity. Biochem Biophys Res Commun. 2015;463(4):1210–7. doi: 10.1016/j.bbrc.2015.06.086. PMID: 26086101.

40. Li X, Thompson D, Kumar B, DeMartino GN. Molecular and cellular roles of PI31 (PSMF1) protein in regulation of proteasome function. J Biol Chem. 2014 ;289(25):17392–405. PMID: 24770418; PMCID: PMC4067172.

41. Michael ARM, Amaral BC, Ball KL, et al. Cell fixation improves performance of in situ crosslinking mass spectrometry while preserving cellular ultrastructure. Nat Commun 2024; 15, 8537. 10.1038/s41467-024-52844-y.

42. Grice GL, Lobb IT, Weekes MP, et al. The Proteasome Distinguishes between Heterotypic and Homotypic Lysine-11-Linked Polyubiquitin Chains. Cell Rep. 2015;12(4):545–53. PMID: 26190103; PMCID: PMC4533228.

43. Isakov E, Stanhill A. Stalled proteasomes are directly relieved by P97 recruitment. J Biol Chem. 2011;286(35):30274–30283. doi: 10.1074/jbc.M111.240309. PMID: 21733848; PMCID: PMC3162386.

44. Ashworth A, Lord CJ. Synthetic lethal therapies for cancer: what’s next after PARP inhibitors? Nat Rev Clin Oncol. 2018;15(9):564–576. doi: 10.1038/s41571-018-0055-6. PMID: 29955114.

45. Ri M. Endoplasmic-reticulum stress pathway-associated mechanisms of action of proteasome inhibitors in multiple myeloma. Int J Hematol. 2016;104(3):273–80. doi: 10.1007/s12185-016-2016-0.

46. Meyer H, Bug M, Bremer S. Emerging functions of the VCP/p97 AAA-ATPase in the ubiquitin system. Nat Cell Biol. 2012;14(2):117–23. doi: 10.1038/ncb2407. PMID: 22298039.

47. Van Den Boom J, Meyer H. VCP/p97-Mediated unfolding as a principle in protein homeostasis and signaling. Mol. Cell 2018; 69, 182–94.

48. Nishimura N, Radwan MO, Amano M, et al. Novel p97/VCP inhibitor induces endoplasmic reticulum stress and apoptosis in both bortezomib-sensitive and - resistant multiple myeloma cells. Cancer Sci. 2019;110(10):3275–3287. doi: 10.1111/cas.14154. PMID: 31368616; PMCID: PMC6778635.

49. Auner HW, Moody AM, Ward TH, et al. Combined inhibition of p97 and the proteasome causes lethal disruption of the secretory apparatus in multiple myeloma cells. PLoS ONE. 2013;8(9):e74415. doi:10.1371/journal.pone.0074415.

50. Roux B, Vaganay C, Vargas JD, et al. Targeting acute myeloid leukemia dependency on VCP-mediated DNA repair through a selective second-generation small-molecule inhibitor. Sci Transl Med. 2021 13(587): eabg1168. doi: 10.1126/scitranslmed.abg1168. PMID: 33790022.

51. Fiskus WC, Das K, Mill CP, et al. Efficacy of Vcp/p97 Inhibitor, CB-5339, alone and in combinations against high-risk AML, including those with genetic lesion in TP53. Blood. 2022;140 (Supplement 1):8807–8. doi:10.1182/blood-2022-165012.

52. Kilgas S, Ramadan K. Inhibitors of the ATPase p97/VCP: From basic research to clinical applications. Cell Chem Biol. 2023;30:3–21. doi: 10.1016/j.chembiol.2022.12.007. PMID: 36640759.

53. Chou TC. Drug combination studies and their synergy quantification using the Chou-Talalay method. Cancer Res. 2010;70(2):440–6. doi: 10.1158/0008-5472.CAN-09-1947. PMID: 20068163.

54. Beskow A, Grimberg KB, Bott LC, et al. A conserved unfoldase activity for the p97 AAA-ATPase in proteasomal degradation. *Journal of Mol*. Biol. 2009;394(4):732–46.

55. Yamanaka K, Sasagawa Y, Ogura T. Recent advances in p97/VCP/Cdc48 cellular functions. BBA -Molecular Cell Research. 2012;1823(1):130–7.

56. Ye Y, Shibata Y, Kikkert M, et al. Recruitment of the p97 ATPase and ubiquitin ligases to the site of retrotranslocation at the endoplasmic reticulum membrane. Proc Natl Acad Sci U S A. 2005;102:14132–8. doi: 10.1073/pnas.0505006102. PMID: 16186510.

57. Le Moigne R, Aftab BT, Djakovic S, et al. The p97 inhibitor CB-5083 is a unique disrupter of protein homeostasis in models of multiple myeloma. Molecular cancer therapeutics. 2017;16(11):2375–86.

58. Brown MG, Driscoll J, Monaco JJ. Structural and serological similarity of MHC-linked LMP and proteasome (multicatalytic proteinase) complexes. Nature. 1991;353(6342):355–7. doi: 10.1038/353355a0.

59. Rana PS, Ignatz-Hoover JJ, Kim B-G, et al, HDAC6 inhibition releases HR23B to activate proteasomes, expand the tumor immunopeptidome and amplify T-cell antimyeloma activity. Cancer Research Communications, 2024; 4(6): 1517–32.

60. Rana PS, Ignatz-Hoover JJ, Guo C, et al., Immunoproteasome Activation Expands the MHC Class I Immunopeptidome, Unmasks Neoantigens, and Enhances T-cell Anti-Myeloma Activity. Mol Cancer Ther. 2024;23(12):1743–60. PMID: 39210605; PMCID: PMC11612626.

61. Gu X, Nardone C, Kamitaki N, Mao A, Elledge SJ, Greenberg ME. The midnolin-proteasome pathway catches proteins for ubiquitination-independent degradation. Science. 2023;381(6660):eadh5021. doi: 10.1126/science.adh5021. PMID: 37616343;

62. Richardson PG, Sonneveld P, Schuster MW, et al.. Bortezomib or high-dose dexamethasone for relapsed multiple myeloma. New England journal of medicine. 2005;352(24):2487–98.

63. Richardson PG, Sonneveld P, Schuster M, et al.. Extended follow-up of a phase 3 trial in relapsed multiple myeloma: final time-to-event results of the APEX trial. Blood. 2007;110(10):3557–60. doi: 10.1182/blood-2006-08-036947.

