## Supplementary for "Synthetic lethality of p97 inhibition and PSMC2-overexpression overcomes bortezomib resistance in multiple myeloma"

**APEX Trial Data Analysis**

International, prospective, randomized, phase III study conducted worldwide that enrolled 669 patients with relapsed and/or refractory MM (RRMM) treated with 1-3 lines of prior therapy. Patients were randomized to receive either bortezomib (327 patients) or high-dose dexamethasone (standard therapy, 330 patients) and consented to genomic analyses of pretreatment tumor samples. Microarray data was analyzed from BM CD138+ cells obtained at enrollment.

*Identify Proteasome Genes that correlate with: Response to Btz (but not Dex): PSMD7, PSMA3, PSMD1, PSMD12, PSMC6, PSMB5, PSMB4,* ***PSMC2****, PSMA4, PSMD11, PSMD13 and PSMD9.*

*Identify Proteasome Genes Correlate with OS following treatment with Btz:*
***PSMC2****, PSMB4, PSMC6, PSME4.*

**CoMMpass Study**

Observational study of NDMM. MD choice of treatment. Blood and BM collected on enrollment and analyzed by NGS to define DNA mutations and copy number, RNA expression.

*Proteasome Genes that Correlate with: Duration of Response: PSMC1, PSMC2*

*Proteasome Genes that Correlate with PFS: PSMC1,* ***PSMC2****, PSMC3, PSMC5 and PSMC6.*

*Proteasome Genes that Correlate OS: PSMC1,* ***PSMC2,*** *PSMC3, PSMC5, PSMC6.*

**Biochemical, Cellular, Genetic and Pharamcologic Studies**

- Engineer *PSMC2 OE and KD* in MMCLs and determine the effect on proteasome activity, PI sensitivity, ER stress response and UPR.
- Synthetic lethality pharmacologic screen to identify drugs that selectively kill *PSMC2* OE cells.
- Determine synthetic lethality and drug synergism between Btz and p97/VCP AAA-ATPase inhibitors in MMCLs, patient CD138$^{+}$ cells and xenograft humanized mouse.

**Figure S1.** Overview of the methods and experiments used in the present study to analyze he APEX and CoMMpass datasets.

**Kaplan-Meier PFS Curves Following Treatment with Bortezomib**

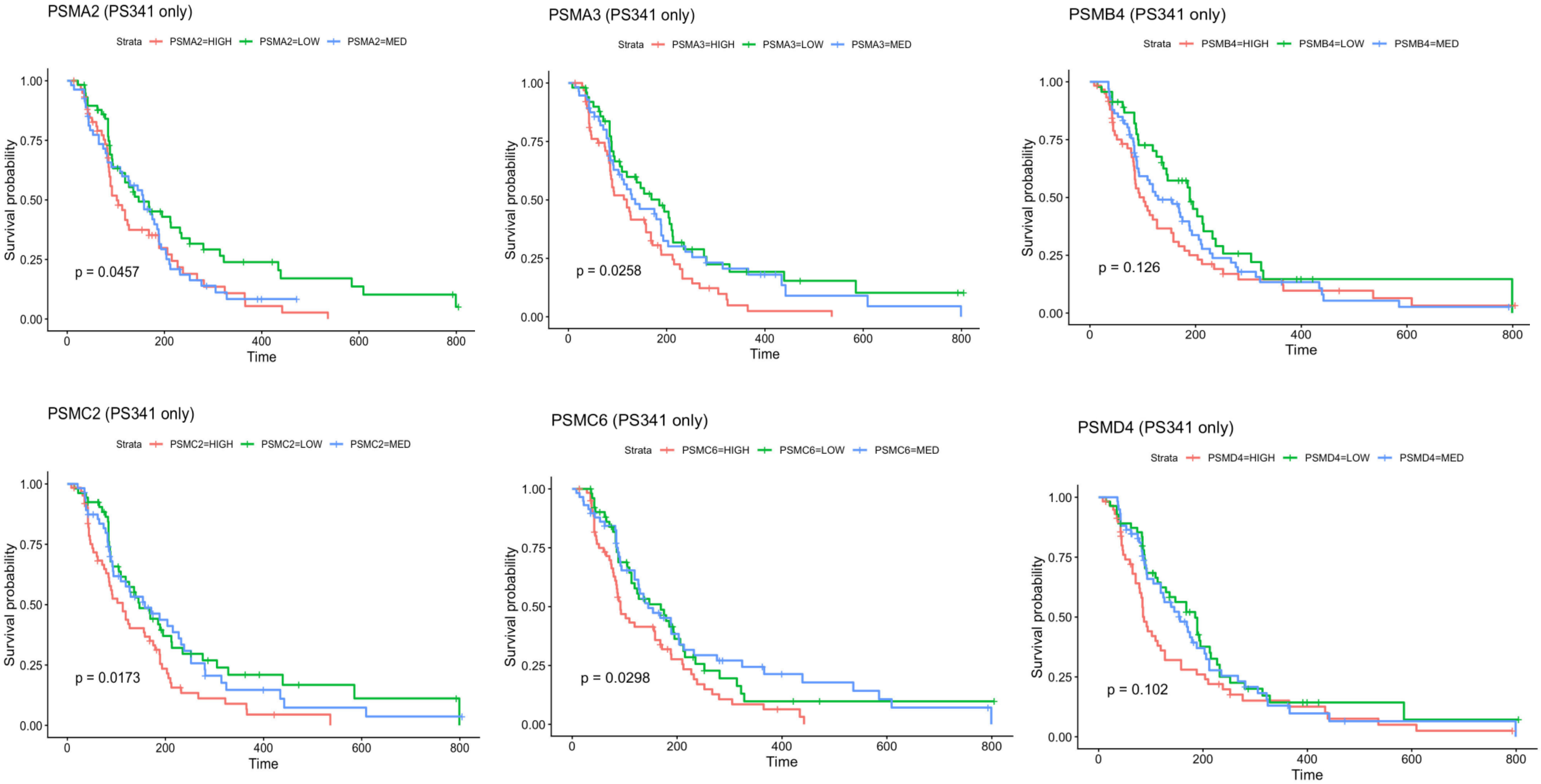


**Figure S2**. Proteasomes genes that correlated with increased risk of inferior PFS following Btz treatment. However, greater expression of *PSMB4* also correlated with inferior PFS risk after Dex treatment.

### Kaplan-Meier PFS Curves for Following Treatment with Dexamethasone

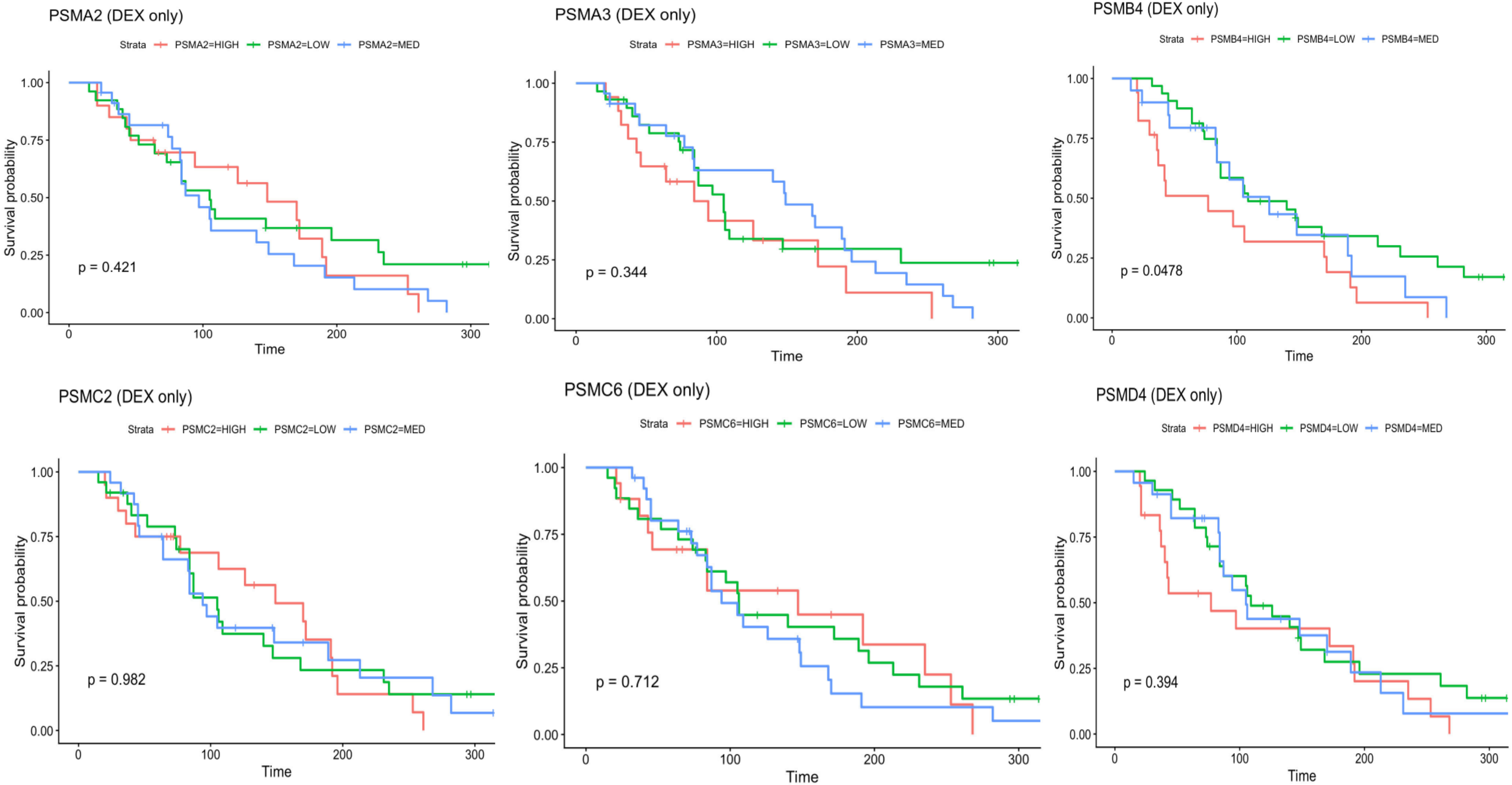


**Figure S3**. Shown is the correlation of proteasomes genes that correlated with increased risk of inferior PFS following Dex treatment. Greater expression of *PSMB4* also correlated with inferior PFS risk after Dex treatment.

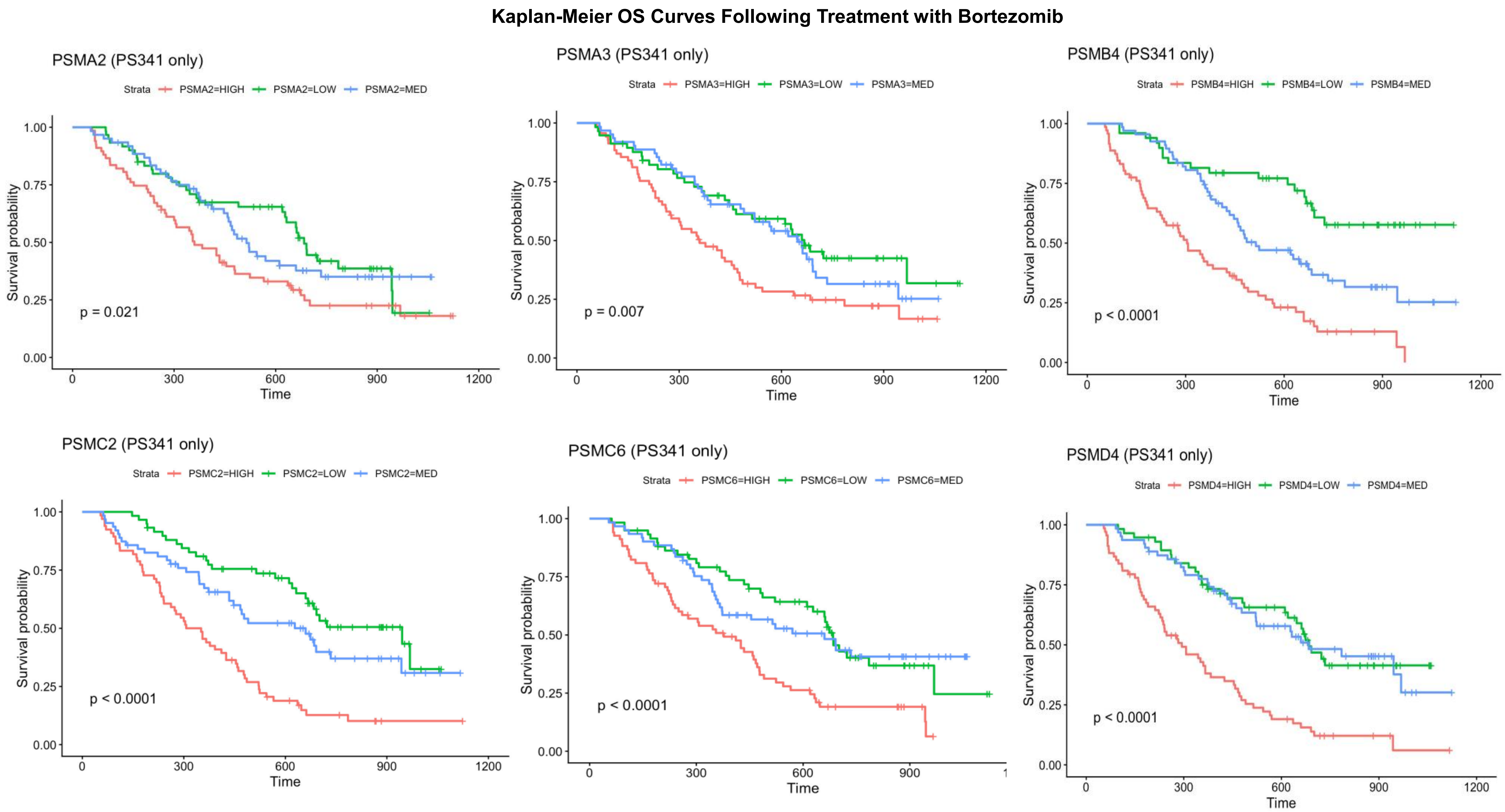


**Figure S4**. Proteasomes genes that correlated with increased risk of inferior OS following Btz treatment. Greater expression of *PSMA2, PSMA3*, and *PSMC6* also correlated with inferior OS in response to Dex treatment.

**Figure S5**

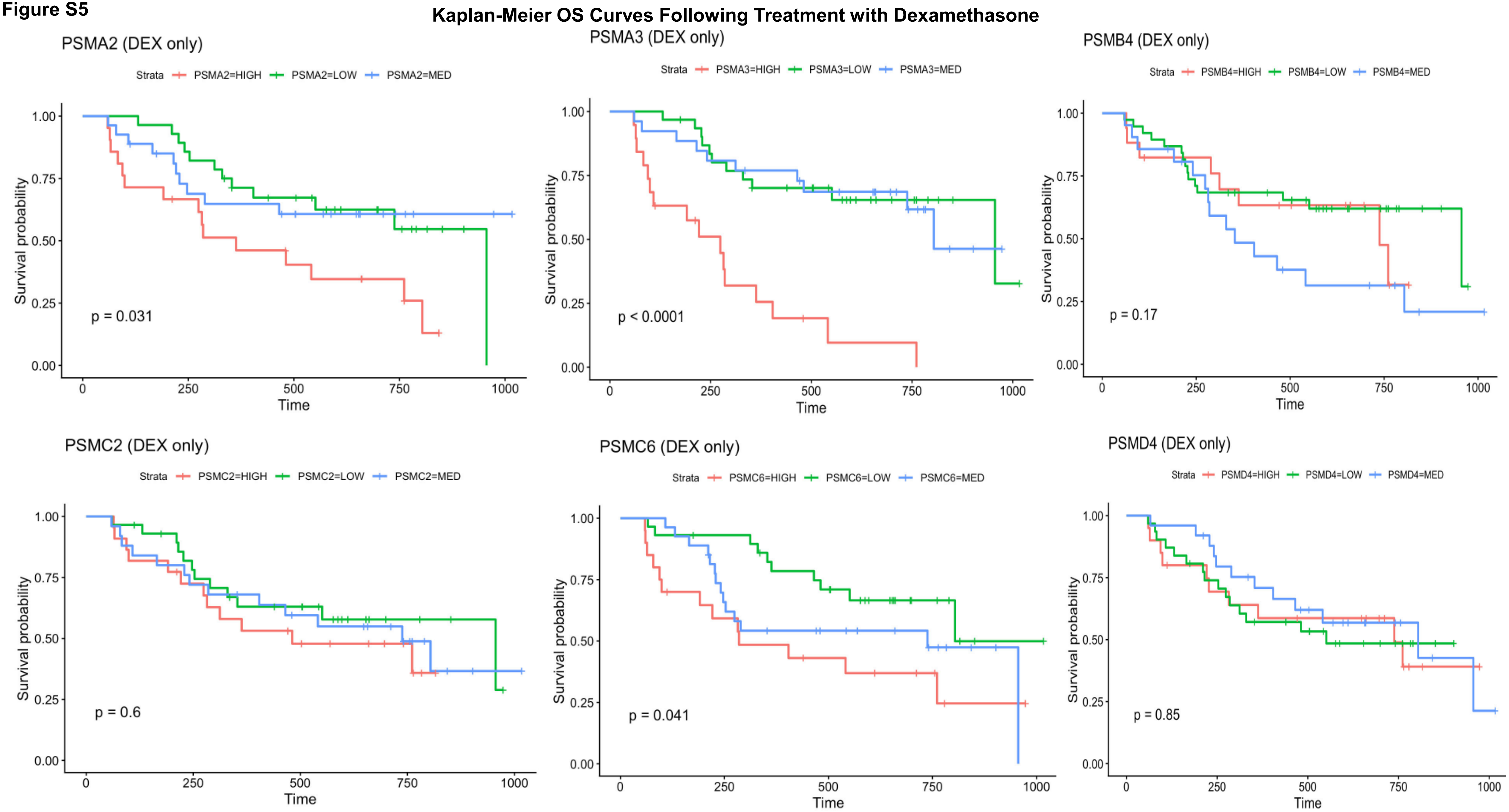


**Figure S5**. Proteasomes genes that correlated with increased risk of inferior OS following Dex treatment. Greater expression of *PSMA2, PSMA3*, and *PSMC6* also correlated with inferior OS in response to Dex.

### Kaplan-Meier PFS Curves Following All Treatments in CoMMpass

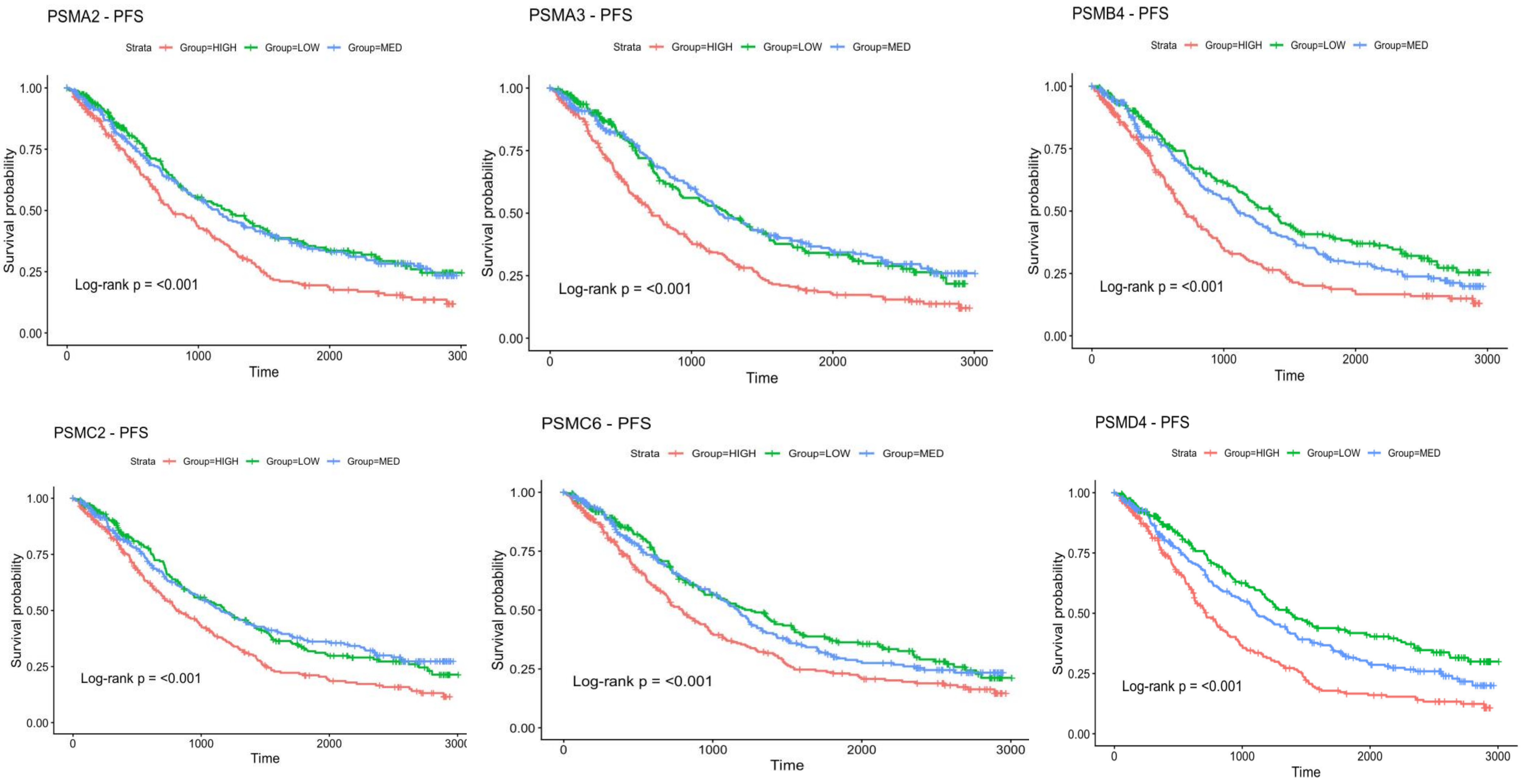


**Figure S6**. Proteasomes genes that correlated with increased risk of inferior PFS in CoMMpass following all treatments.

**Kaplan-Meier OS Curves Following All Treatments in CoMMpass**

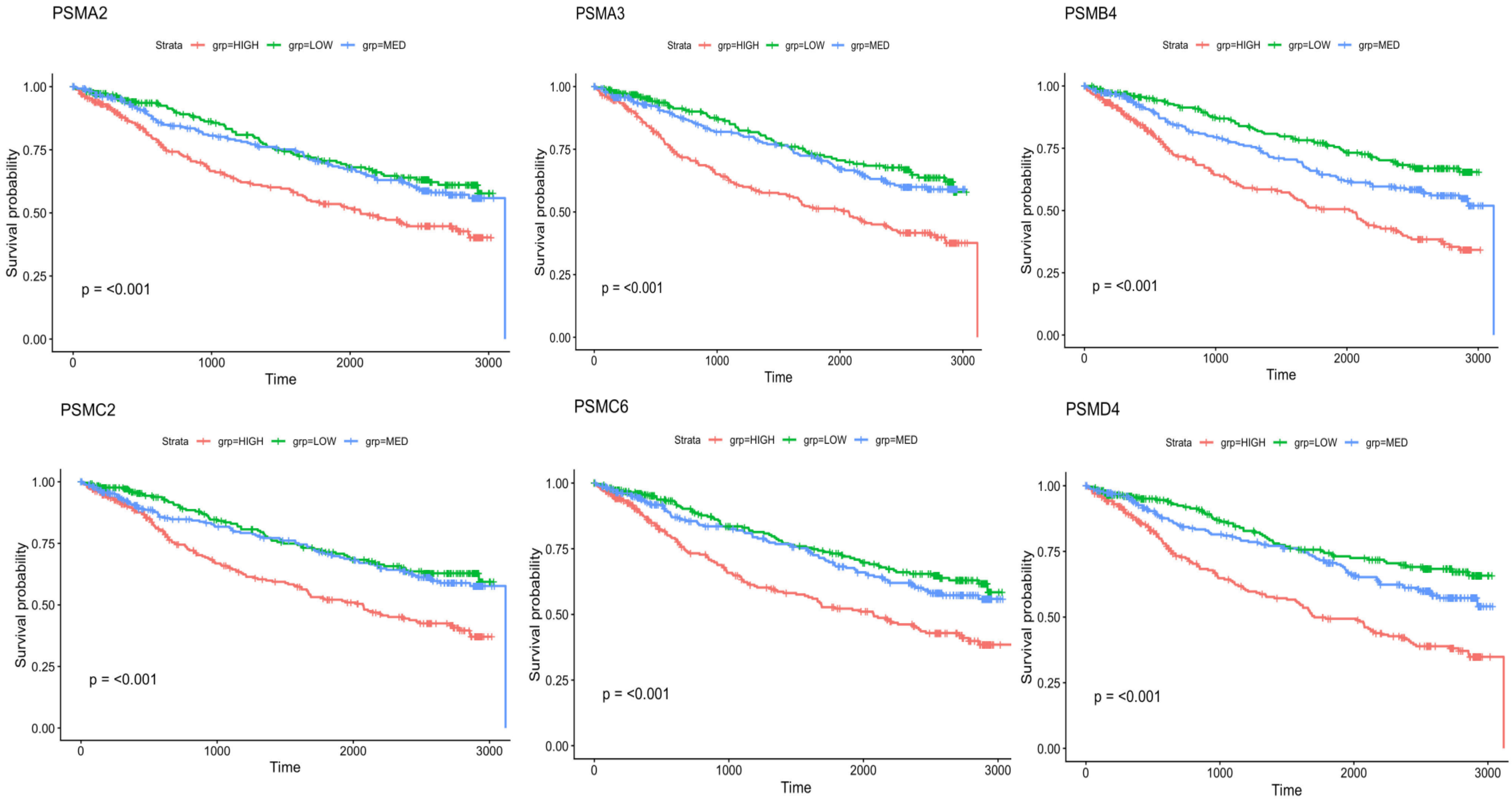


**Figure S7**. Proteasomes genes that correlated with increased risk of inferior OS in CoMMpass following all treatments.

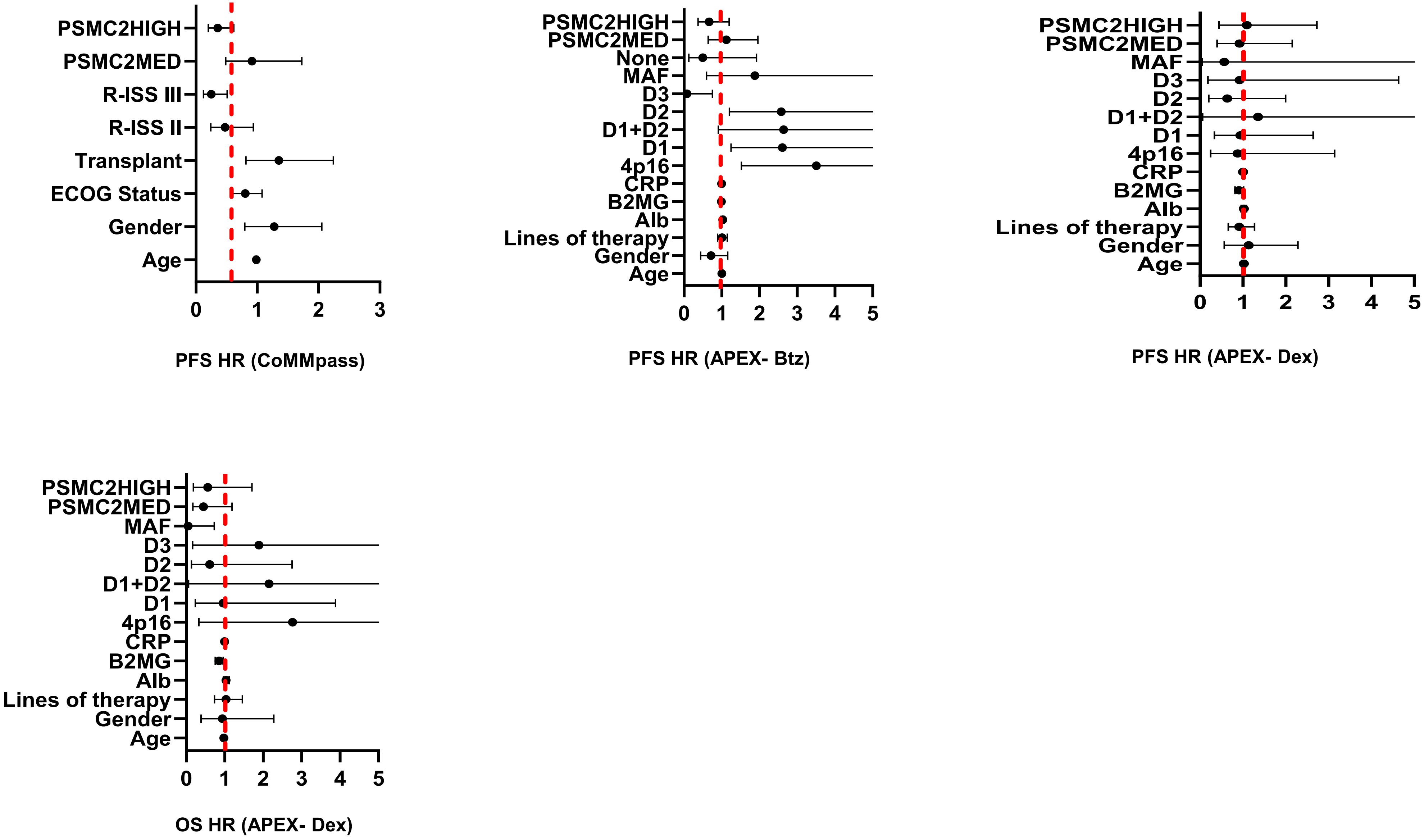


**Fig. S8. Multivariate analysis of high *PSMC2* expression in the APEX an CoMMpass datasets**. Shown is the correlation of high *PSMC2* expression with PFS in the CoMMpass study and PFS for patients treated with either Btz or Dex in the APEX study. High *PSMC2* expression did not correlate with PFS or OS in the APEX study after treatment with Dex. High *PSMC2* expression correlated with inferior OS in both the APEX and CoMMpass studies (Fig. 1h).

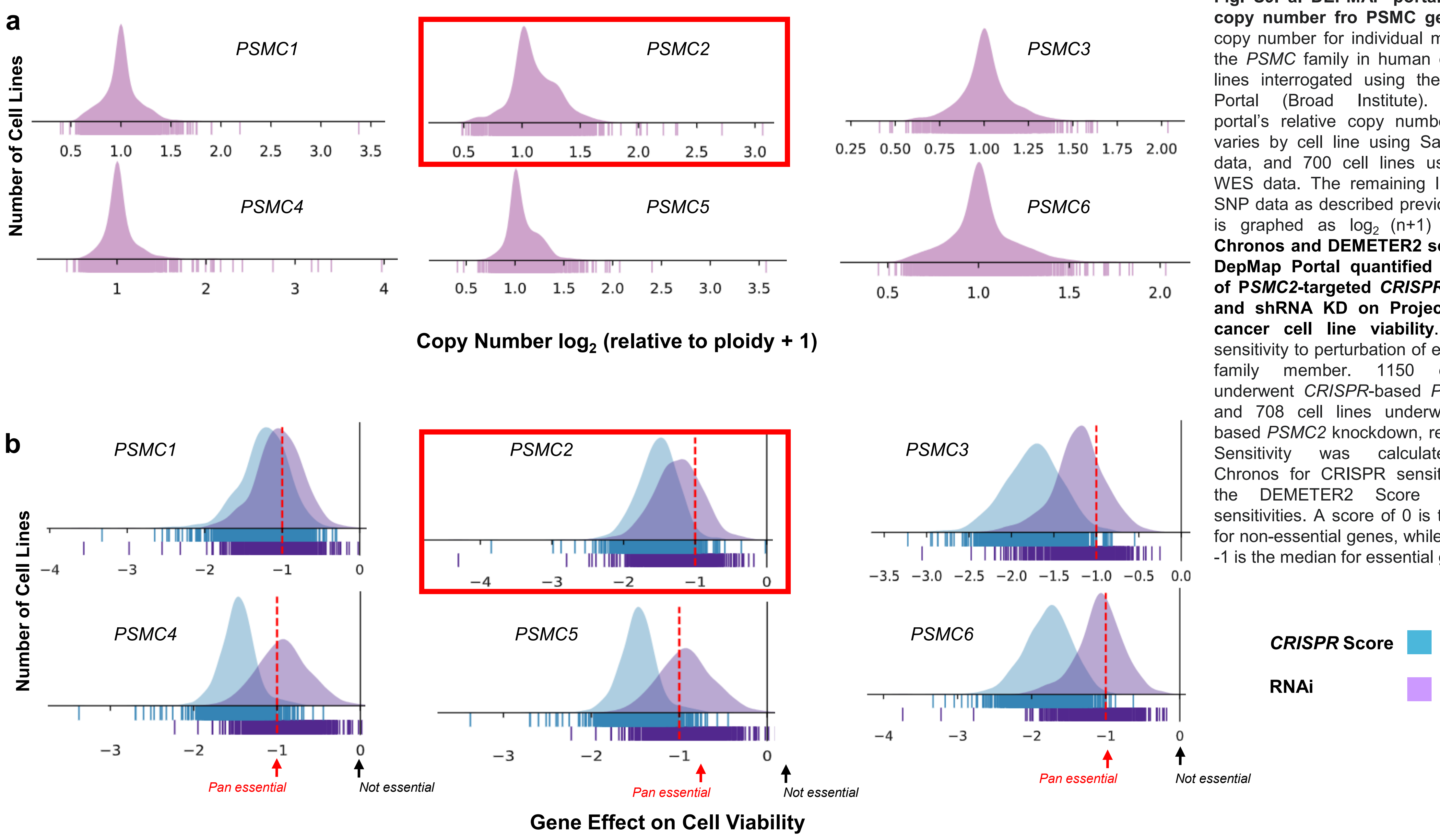


**Fig. S9. a. DEPMAP portal's relative copy number fro PSMC genes.** DNA copy number for individual members of the *PSMC* family in human cancer cell lines interrogated using the DEPMAP Portal (Broad Institute). DEPMAP portal's relative copy number pipeline varies by cell line using Sanger WES data, and 700 cell lines using Broad WES data. The remaining lines utilize SNP data as described previously. Data is graphed as $\log_2$ (n+1) ploidy. **b. Chronos and DEMETER2 scales from DepMap Portal quantified the effect of P*SMC2*-targeted *CRISPR/Cas9* KO and shRNA KD on Project Achilles cancer cell line viability.** Cell line sensitivity to perturbation of each *PSMC* family member. 1150 cell lines underwent *CRISPR*-based *PSMC2* KO and 708 cell lines underwent RNAi-based *PSMC2* knockdown, respectively. Sensitivity was calculated using Chronos for CRISPR sensitivities and the DEMETER2 Score for RNAi sensitivities. A score of 0 is the median for non-essential genes, while a score of -1 is the median for essential genes.

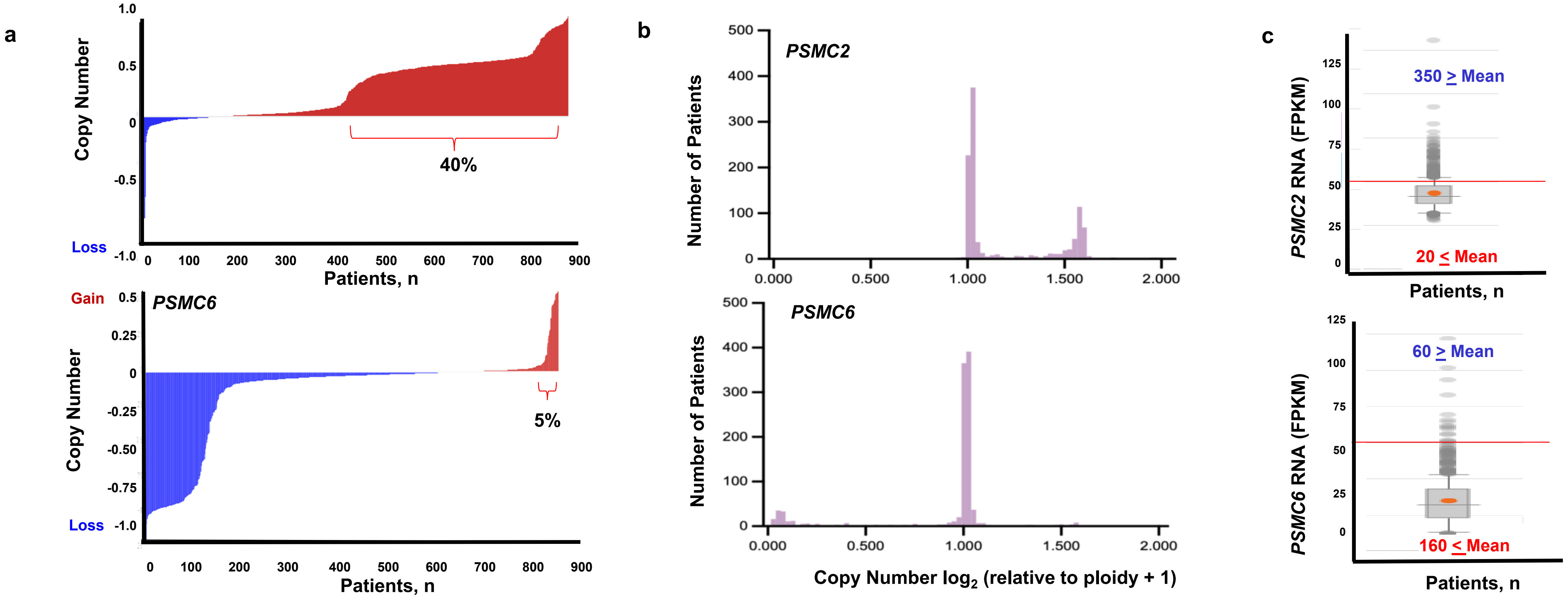


**Figure S10.** **a.** Waterfall plot to represent the magnitude of *PSMC2* and *PSMC6* copy number alteration (Y-axis) in CD138$^{+}$ cells from individual patients enrolled on the CoMMpass study (n=1077). Patients with increased *PSMC* copy number are indicated in red and patients with decreased *PSMC* copy number are indicated in blue (X-axis). **b.** Number of patients (Y-axis) vs. copy number $\log_2$ (relative to ploidy + 1) for *PSMC2* or *PSMC6* using the CoMMpass dataset. **c.** Box and whisker plots of RNA expression levels of *PSMC2* and *PSMC6* from the CoMMpass study. Of note, ~350 patients exhibit RNA levels above the mean for *PSMC2* (n = 420) while 60 patients exhibit *PSMC6* levels above the mean (n = 220).

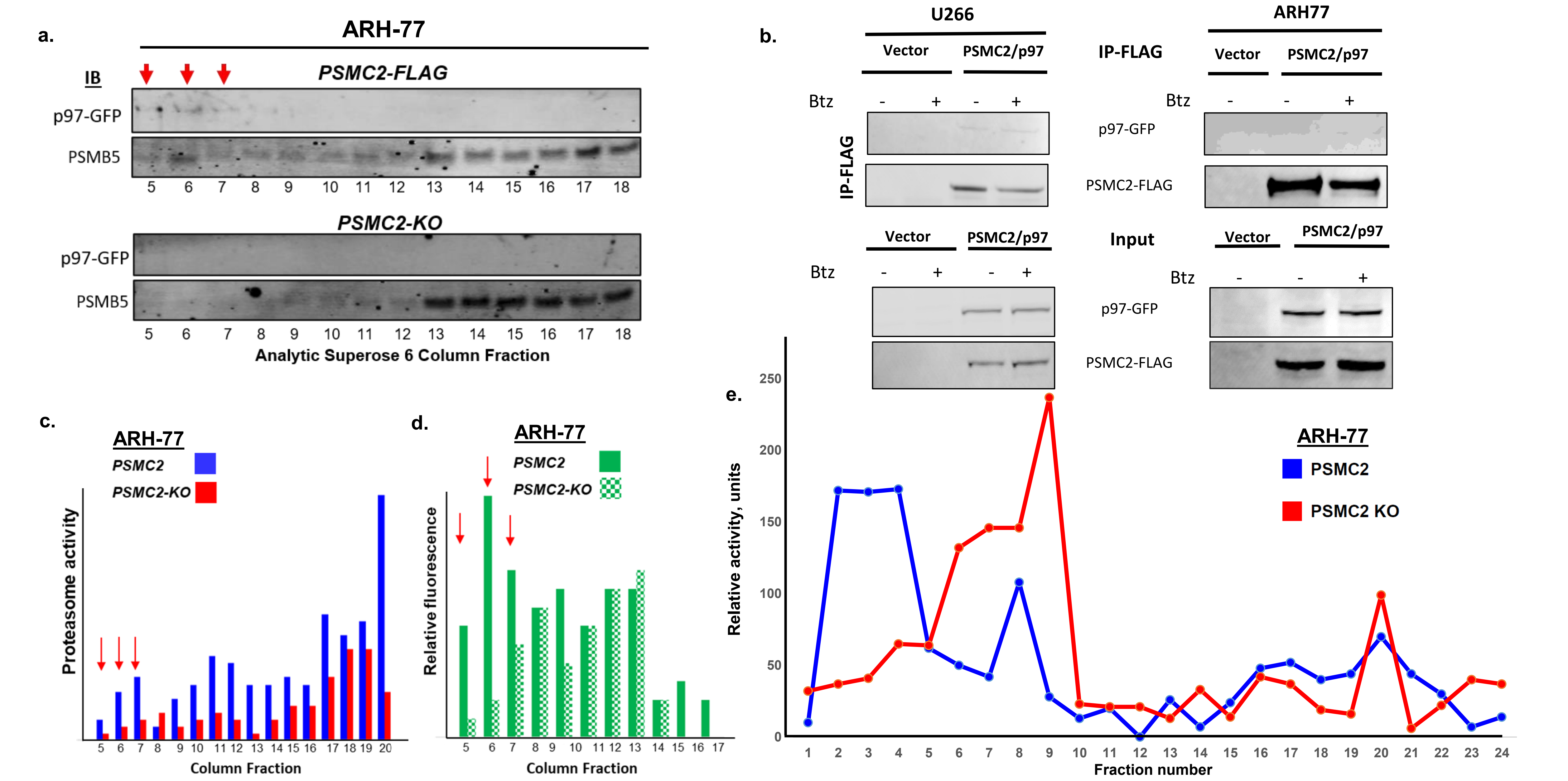


**Figure S11. a.** Western blot of gel filtration fractions from ARH-77 cell lysates for p97-GFP and PSMB5. p97-GFP is detected in the 26S proteasome fractions (red arrows) from ARH-77 cells that express *PSMC2-FLAG* but not *PSMC2-KO* cells. **b**. Western blots for p97-GFP and PSMC2-FLAG from co-immunoprecipitates of U266 and ARH-77 cells. Btz treatment increases p97-GFP association with PSMC2-FLAG. **c**. Proteasome hydrolysis of Suc-LLVY-mca assayed in Superose 6 column fractions from affinity-purified proteasomes obtained from *PSMC2-expressing* and *KO* lysates. Red arrows indicate the peaks associated with 26S and 20S complexes, respectively. **d.** Fluorescence measured at Ex $488_{nm}$/Em $510_{nm}$ in column fractions from affinity-purified proteasomes obtained from *PSMC2*-expressing and *KO* cell lysates. **e.** Fluorescence measured at Ex $488_{nm}$/Em $510_{nm}$ in SEC column fractions from affinity-purified proteasomes obtained from *PSMC2*-expressing cell lysates with and without Btz treatment.

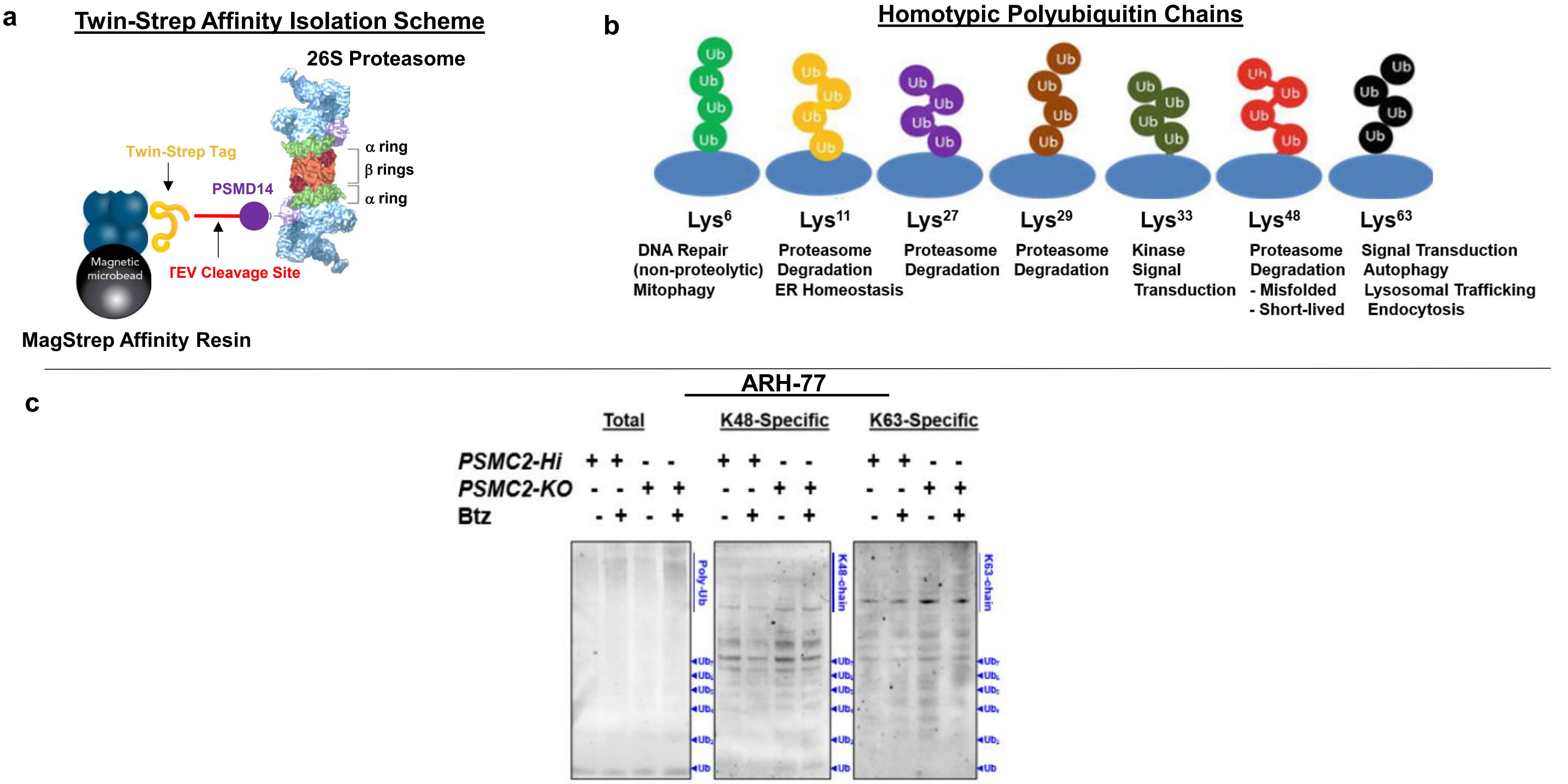


**Figure S12. a.** Twin-Strep affinity isolation scheme to affinity purify 26S proteasomes from MM cells**. b.** Homotypic polyubiquitin chains are polymer structures where ubiquitin monomers link together uniformly through the exact same single lysine residue (K6, K11, K27, K29, K33, K48, or K63). Distinct canonical homotypic chain types forming unique 3-D structures that signal specific cellular fates. **c.** Western blot of homotyp ubiquitin chains in ARH-77 cell lysates. Cells were treated with cross-linker, treated as indicated with Btz, and proteasomes affinity purified. Puriifed proteisn were then separated by SDS-PAGE and probed using indicated antibodies.

**a. Expression of *PSMC2* in 1080 Human Cancer Cell Lines**

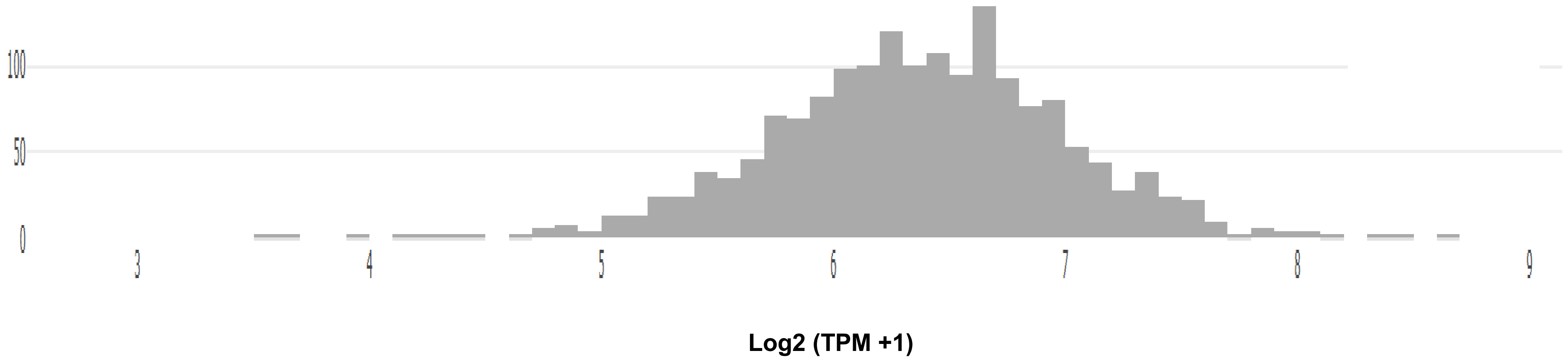


**b. Expression of *PSMC2* in 188 Lymphoid Cancer Cell Lines**

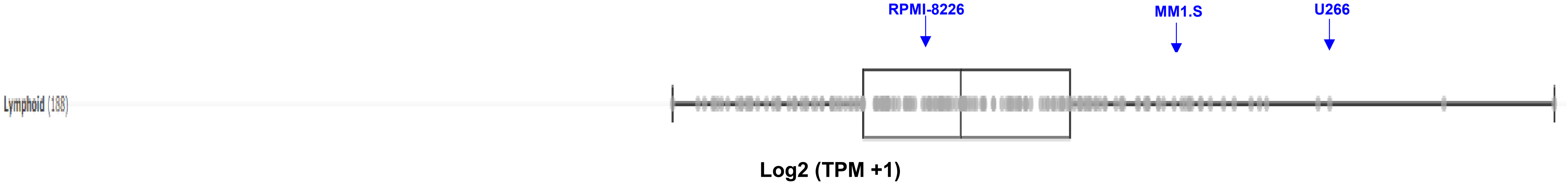


**Figure S13. a.** Shown is the relative expression of *PSMC2* generated from the Cancer Dependency Map for all cancer cell lines (DepMap portal, Broad Institute). **b.** Shown is the relative expression of *PSMC2* generated from the Cancer Dependency Map for lymphoid (including multiple myeloma) cell lines.

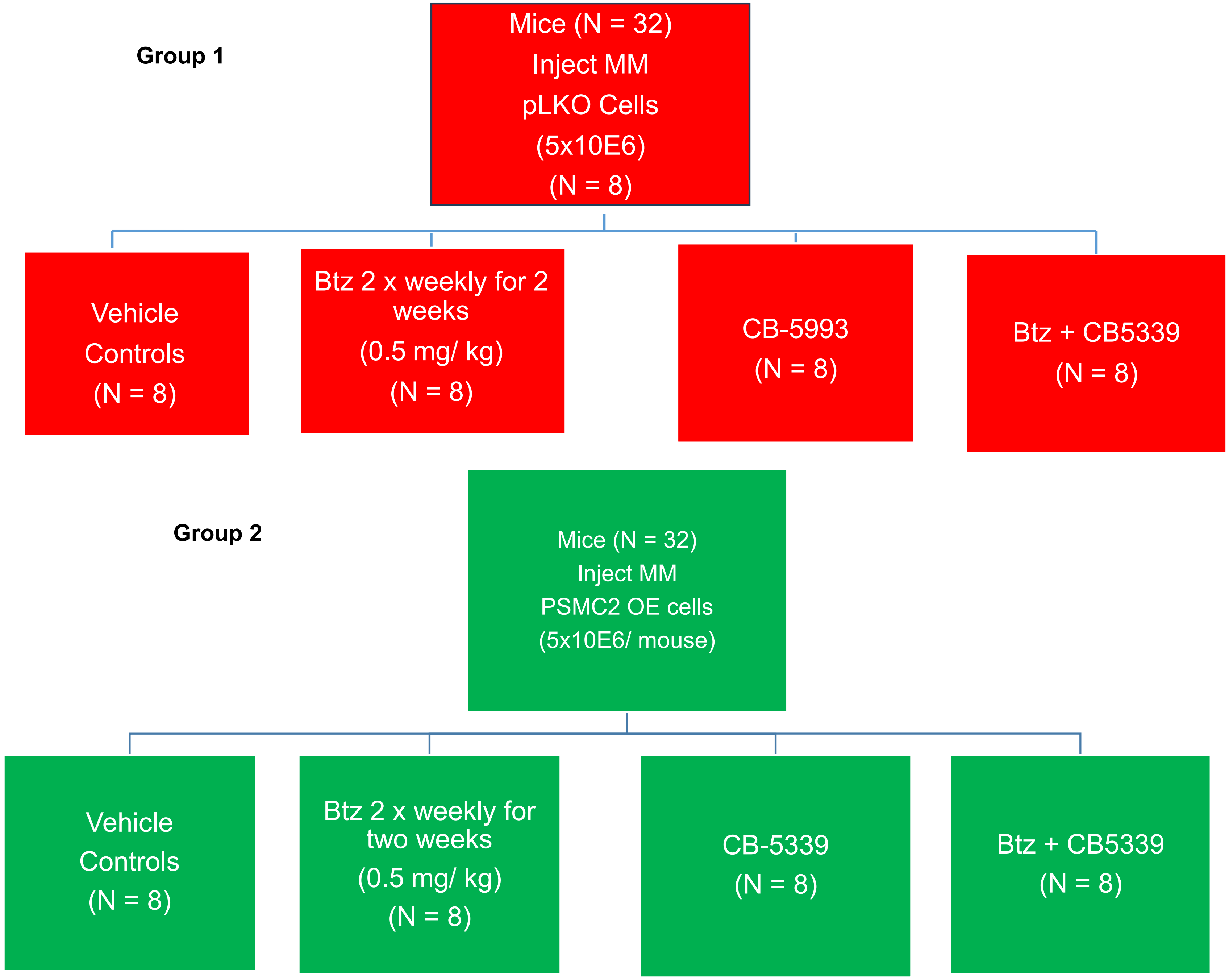


**Figure S14.** Cohort Diagram for NSG Mouse Studies on the effect of the p97 ATPase activity inhibitor CB5339 +/- Btz.

Figure S15

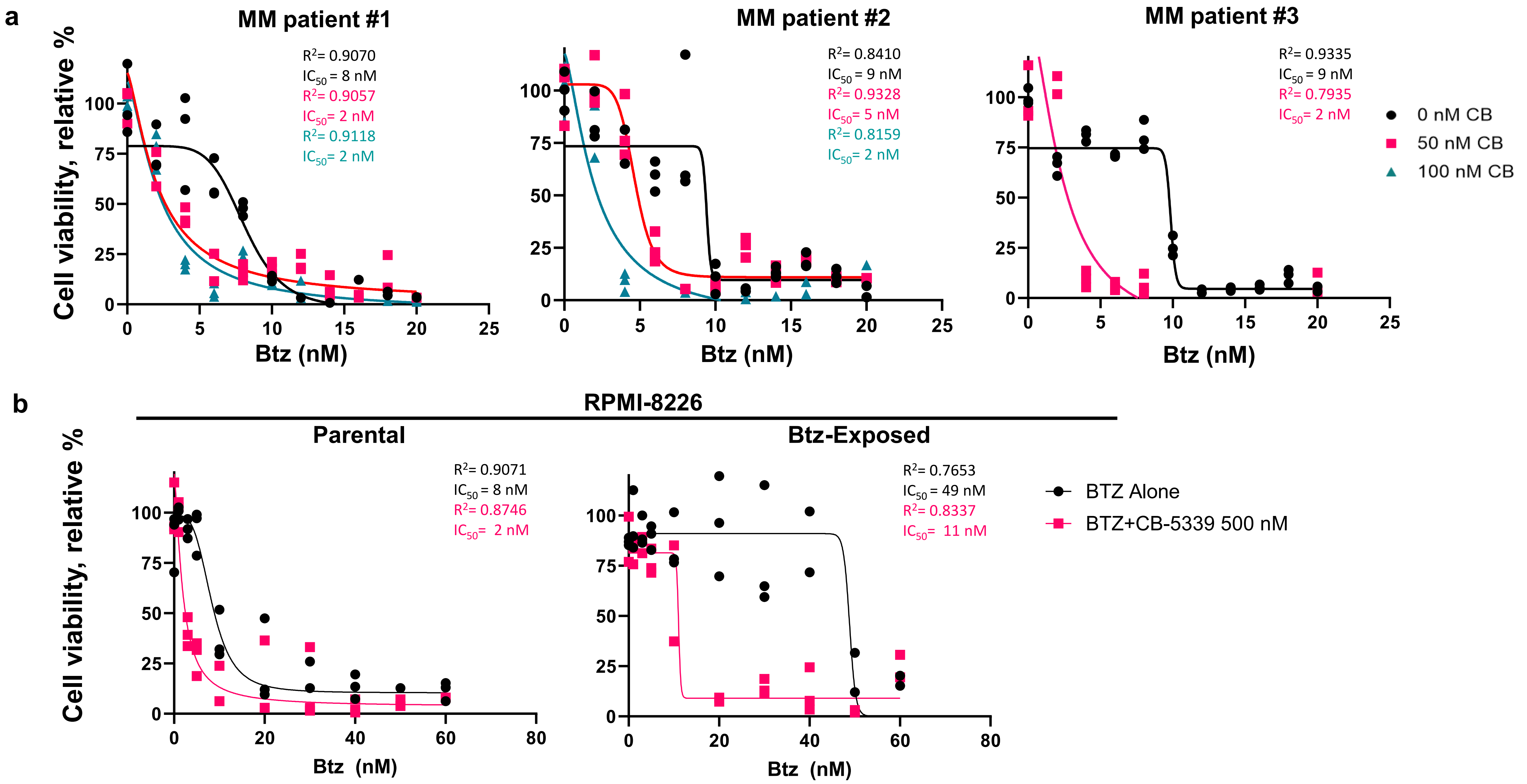


**Fig. S15. a.** Effect of Btz alone and combined with CB-5339 on the viability of MM cells from drug exposed patients. **b.** Effect of Btz alone and combined with CB-5339 on the viability of parental and Btz-resistant RPMI8226 cells. Viability was determined using the XTT assay. All bioassays were performed in triplicate.

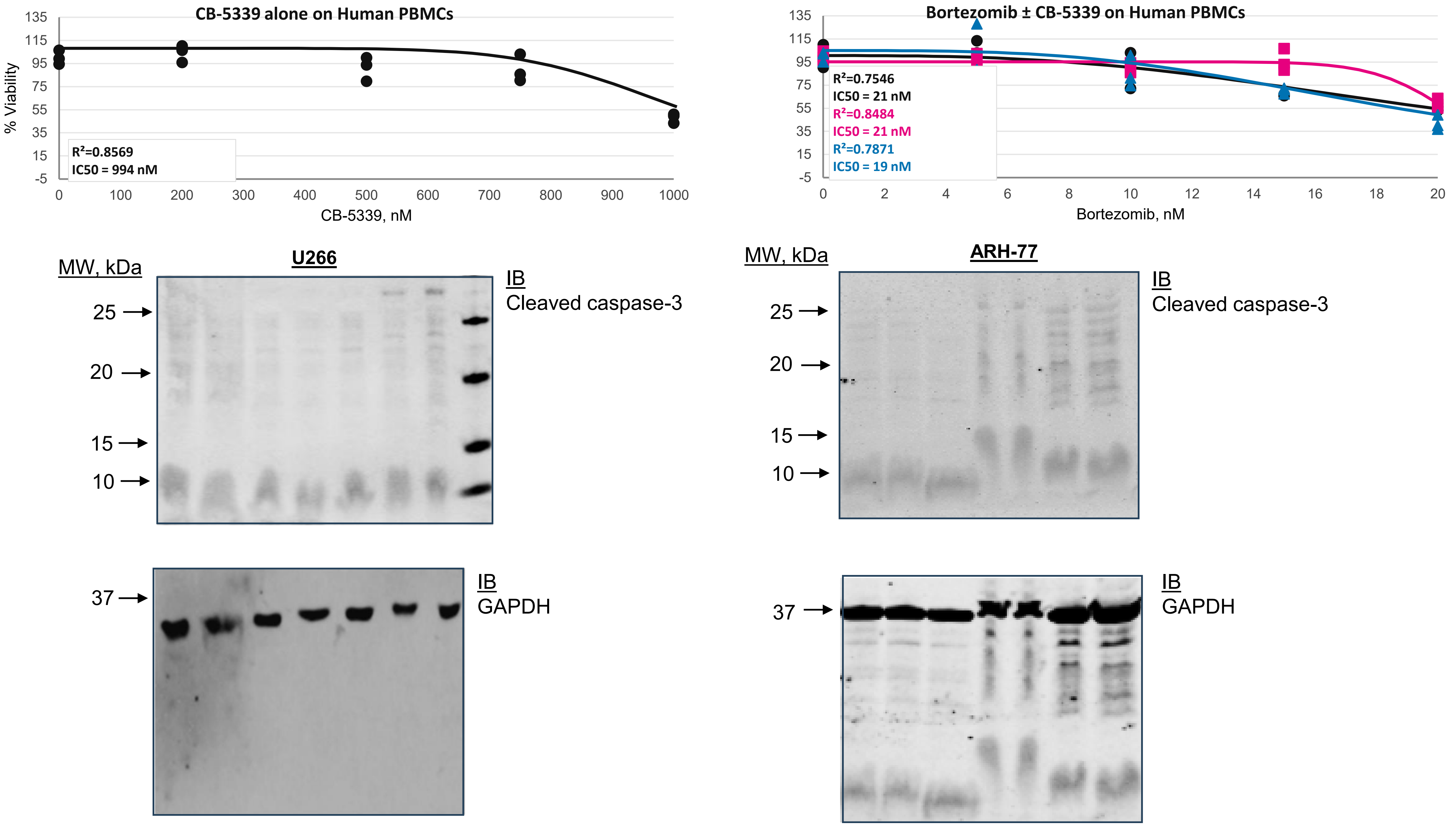


**Fig. S16. a.** Effect of Btz alone and combined with CB-5339 on the viability of healthy PBMCs. **b.** Western blot of U266 and ARH-77 cells treated with CB-5339 and Btz at the indicated concentrations. Lysates were prepared and probed using an antibody to cleaved caspase-3.
